# Characterization of the *Cannabis sativa* glandular trichome epigenome

**DOI:** 10.1101/2024.07.04.602151

**Authors:** Lee J. Conneely, Bhavna Hurgobin, Sophia Ng, Muluneh Tamiru-Oli, Mathew G. Lewsey

**Affiliations:** La Trobe Institute for Sustainable Agriculture and Food, La Trobe University, AgriBio Building, Bundoora, VIC 3086, Australia; Australian Research Council Research Hub for Medicinal Agriculture, La Trobe University, AgriBio Building, Bundoora, VIC 3086, Australia; Australian Research Council Centre of Excellence in Plants for Space, La Trobe University, Bundoora, Victoria, Australia

**Keywords:** Cannabis sativa, Glandular trichomes, Specialised metabolism, Multiomics, H3K4me3, H3K27me3, H3K56ac, H2A.Z, Chromatin immunoprecipitation, Gene Regulation, Cis-regulatory element

## Abstract

**Background:** The relationship between epigenomics and plant specialised metabolism remains largely unexplored despite the fundamental importance of epigenomics in gene regulation and, potentially, yield of products of plant specialised metabolic pathways. The glandular trichomes of *Cannabis sativa* are an emerging model system that produce large quantities of cannabinoid and terpenoid specialised metabolites with known medicinal and commercial value. To address the lack of epigenomic data in plant specialised metabolism, glandular trichomes, and *C. sativa*, we mapped H3K4 trimethylation, H3K56 acetylation, H3K27 trimethylation post-translational modifications and the histone variant H2A.Z, using chromatin immunoprecipitation, in glandular trichomes, leaf, and stem tissues. Corresponding transcriptomic (RNA-seq) datasets were integrated, and tissue-specific analyses conducted to relate chromatin states to glandular trichome specific gene expression.

**Results:** Cannabinoid and terpenoid biosynthetic genes, specialised metabolite transporters, and defence related genes, were co-located with distal H3K56ac chromatin, a histone mark that flanks active distal enhancers *in planta*, exclusively in glandular trichomes. Glandular trichome specific H3K4 trimethylated chromatin was associated with genes involved in specialised metabolism and sucrose and starch metabolism. Bi-valent chromatin loci specific to glandular trichomes, marked with H3K4 trimethylation and H3K27 trimethylation, was associated with genes of MAPK signalling pathways and plant specialised metabolism pathways, supporting recent hypotheses that implicate bi-valent chromatin in plant defence. The histone variant H2A.Z was largely found in intergenic regions and enriched in chromatin that contained genes involved in DNA homeostasis.

**Conclusion:** We report the first genome-wide histone post-translational modification maps for *C. sativa* glandular trichomes, and more broadly for glandular trichomes in plants. Our findings have implications in plant adaptation and stress response and provide a basis for enhancer-mediated, targeted, gene transformation studies in plant glandular trichomes.

## Background

Histone post-translational modifications and histone variants are features of plant epigenomes that influence DNA templated activities including gene regulation and, through this, tissue and cell function [1–3]. H3K4 trimethylation (H3K4me3) and H3K27 trimethylation (H3K27me3) are the most extensively studied histone post-translational modifications in plants and animals [4–6]. H3K4me3 and H3K27me3 have opposing functions on gene expression [7, 8]. H3K4me3 is deposited by trithorax complexes in the 5’ untranslated region (UTR) of actively transcribed genes where it is required for RNA polymerase II pause-release and transcriptional elongation [9–11]. Conversely, H3K27me3 is deposited by polycomb repressor complex 2 (PRC2) and contributes to the formation of facultative heterochromatin, referred to as polycomb chromatin, that is enriched in the gene body of repressed genes. Furthermore, H3K27me3 PRC2 mediated gene repression operates synergistically with polycomb repressor complex 1 (PRC1) in the monoubiquitylation of the histone variant H2A.Z to repress gene expression [12, 13]. Similar to H3K27me3, H2A.Z in plants is enriched in the gene body of repressed genes [14].

H3K4me3 and H3K27me3 have primarily antagonistic effects on gene expression, but they may also co-localise on the same nucleosome in a phenomenon known as bivalency. It is hypothesised that bi-valent chromatin containing antagonistic histone post-translational modifications results in poised states of gene expression, whereby plants can rapidly upregulate and fine tune spatio-temporal gene expression through the integration of various environmental cues [15]. Vernalization is the most well studied example of H3K4me3-H3K27me3 bivalency in plants, whereby temperatures influence the ratio of H3K4me3 to H3K27me3 at a bi-valent locus associated with the floral development repressor gene *FLC* [16].

This modulates *FLC* expression through gene silencing, allowing floral development when temperature increases in spring. More recently the balance between facultative heterochromatin associated H3K27me3 and euchromatin associated H3K18 acetylation has been proposed to govern expression of genes involved in the biosynthesis of the antimicrobial specialised metabolite camalexin in the presence of the pathogen associated molecular pattern (PAMP) Flg22 [17].

Enhancer elements are short stretches of DNA that contain transcription factor binding motifs of approximately 7-22 bp [18]. Interactions between enhancer elements and gene promoters drive gene transcription, many enhancers having tissue and cell-type specific activity [19]. Enhancers can regulate single or multiple gene targets. They can be found within genes, often in introns, where they directly regulate expression of that gene and/or its neighbours, as well as in intergenic regions [20]. Enhancers found in intergenic regions may regulate expression of proximal (≤1.5 Kbp) genes, or distal genes that are located from thousands to a million base pairs away (termed distal enhancer elements) [21–23]. Distal enhancers mediate gene expression via long-range chromatin interactions known as chromatin looping [23, 24]. Enhancers can drive transcription of tissue and cell-type specific genes and thereby influence tissue and cell type identity, making them a useful resource to understand cell type specificity of gene regulation [25, 26].

Chromatin features specific to enhancers and transcription factor binding sites can be used to discover novel regulatory elements [27–29]. In both plants and animals, active enhancers are typically flanked by narrow regions of acetylated euchromatin in the tissue or cell type where they are active, while the transcription factor binding site itself is typically exposed DNA. There is evolutionary conservation of H3K56 acetylation (H3K56ac) flanking distally accessible chromatin regions, correlating positively with the expression of genes local to that region, in at least 13 species of angiosperm including *Sorghum bicolor*, *Zea mays*, and *Oryza sativa* show.

This has led to the proposition that H3K56ac is likely a histone post-translational modification that marks distal enhancer elements in plants [30, 31]. Furthermore, it has been demonstrated that the corresponding mammalian distal enhancer mark H3K27 acetylation (H3K27ac) can be leveraged to mine active transcription factor binding sites [32]. H3K56ac is also enriched in the promoters of transcribed genes similar to H3K4me3, it is thought to play a general role in the affinity for DNA to nucleosomes promoting relaxed euchromatin status [30].

*Cannabis sativa* is a predominantly dioecious, dicotyledonous, plant of the Cannabaceae family that has been cultivated for at least 2700 years for both its long, durable fibres and its medicinal properties [33–35]. The medicinal compounds produced by *C. sativa* are the cannabinoids, a group of species-specific terpenophenolic compounds noted for their medicinal and psychoactive properties [36, 37]. Moreover, the organs that produce specialised metabolites like cannabinoids, the glandular trichomes, have garnered the interest of the plant science community as potential new models for the study of plant cell and tissue development [38]. Most recently there has been interest in potential engineering of glandular trichomes, due to the remarkable quantities of specialised metabolite they are able to produce and sequester away from other plant tissues (e.g. THC 14.98 +/- 2.23% dry weight) [39–42]. However, for glandular trichome engineering to be feasible in *C. sativa* we must understand and be able to manipulate trichome-specific gene expression and specialised metabolism. This would be enabled by mapping epigenomic features, such as histone modifications and variants, and relating them to gene expression characteristics. There is currently a lack of any such data in *C. sativa* or glandular trichomes more broadly.

In this study we generated genome-wide maps of histone post translational modifications H3K4me3, H3K27me3, H3K56ac, and the histone variant H2A.Z for *C. sativa* glandular trichomes, leaves and stems. We made comparisons between tissues to identify trichome specific epigenomic signals. Matched transcriptomic (RNA-seq) datasets were generated for each tissue type to validate histone function in relation to gene expression and to enable functions associated with different epigenome features to be analysed. The relationships of these functions with plant specialised metabolism, abiotic and biotic stress resistance were assessed. Lastly, putative glandular trichome specific enhancer regions were predicted by examining H3K56ac data across tissues and drawing comparisons to differential gene expression analysis.

## Results

### Chromatin immunoprecipitation of histone variants in three *C. sativa tissues*

We first asked if we could generate epigenome wide maps of well characterised histone modifications in *C.* sativa. To this end we mapped the distribution of the epigenomic marks H3K56ac, H3K4me3, H3K27me3, and histone variant H2A.Z across cannabis tissues and relate them to tissue-specific expression of specialized metabolic pathway genes. A chromatin immunoprecipitation sequencing (ChIP-seq) protocol was first optimised for cannabis tissues, as extreme concentrations of specialized metabolites may interfere with analytical methods established in other species (Additional File 1: Fig S1). Immunoprecipitated H3K4me3, H3K27me3, H3K56ac, and histone variant H2A.Z chromatin from glandular trichome, stem (internode), and vegetative leaf tissues were then quantified and processed into next generation sequencing libraries (Additional File 1: Fig. S2, Table S1, Table S2). Next generation sequencing yielded overall good alignment, with the degree of uniquely mapping reads compared with multi-mapping reads highly specific to each histone post-translation modification or histone variant H2A.Z (Additional File 1, Table S3a). Quality control fingerprint plots were consistent with expected distributions for genome-wide narrow peak H3K4me3 and H3K56ac and broad peak H3K27me3 and H2A.Z data, respectively (Additional File 1; Fig. S3). Corresponding RNA-seq libraries were generated in triplicate for each tissue for the purpose of integration with the ChIP-seq data (Additional File 1: Fig. S4, Table S3b).

### Associations of histone post-translational modifications with gene expression in *C. sativa*

We then asked if we could observe well conserved functional properties for these histone modifications with respect to gene expression in *C. sativa*. We assessed the distribution of histone modifications and variants, examining possible associations between the locations of different marks and expressed genes (Fig. 1b-c, Additional File 1: Fig. S5). There were strong positive correlations between read map sites of H3K56ac, H3K4me3, and RNA-seq reads for each tissue, while H3K27me3 and H2A.Z reads tended to negatively correlate with RNA-seq reads (Spearman correlation coefficient, Fig. 1b). Genome-wide visualization of read distributions agreed with this, with actively transcribed genes overlaying reads for both H3K4me3 and H3K56ac (Fig. 1c). The genome-wide overlay between un-transcribed genes, H3K27me3 and H2A.Z was also observed, which may indicate roles in the repression of transcription. A large proportion of reads from H3K27me3 and H2A.Z were from the intergenic regions, with a greater proportion of H3K27me3 reads found in gene features when compared to H2A.Z(Fig. 1d). The majority of H3K4me3 and H3K56ac reads mapped to gene features (Fig. 1d). The distribution of histone marks across genes was consistent with observations in other species [31, 43]. H3K4me3 and H3K56ac were enriched at the transcriptional start site of actively transcribed genes. In other species this thought to facilitate RNA polymerase II pause-release and a relaxed chromatin environment that reduces steric hinderance of trans-acting factors respectively [9, 43, 44]. H3K27me3 and H2A.Z were enriched in the gene bodies of un-transcribed genes, consistent with their synergistic roles in polycomb mediate gene repression [14, 45] (Fig. 1e, Additional File 1: Fig. S6).

**Fig. 1.**
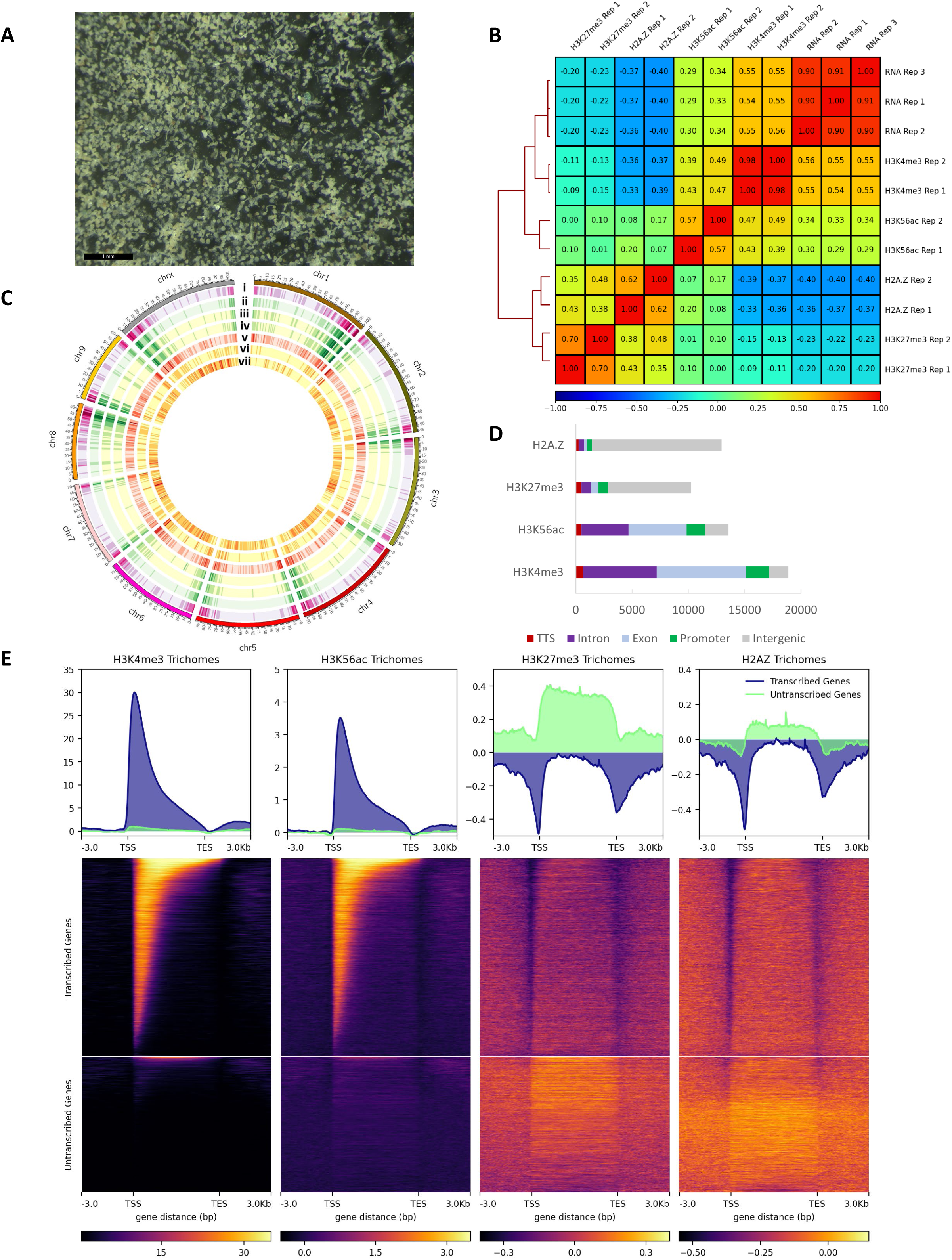
**A** Light microscopy image (1 mm scale bar) of isolated *C.* glandular trichomes. **B** Spearman correlation matrix of RNA-seq and ChIP-seq data for H3K4me3, H3K56ac, H3K27me3, and H2A.Z in glandular trichomes. **C** *C. sativa* karyotype with glandular trichome data density plots (i) all genes (ii) transcribed genes (iii) H3K4me3 peaks (iv) H3K56ac peaks (v) untranscribed genes (vi) H3K27me3 domains and (vii) H2A.Z domains. **D** H2A.Z, H3K27me3, H3K56ac, and H3K4me3 peak annotation. **E** Scale-region gene plots and associated heatmaps of glandular trichome H3K4me3, H3K56ac, H3K27me3, and H2A.Z reads distributed across glandular trichome transcribed genes and untranscribed genes.

### Histone marks co-occur in the *C. sativa* genome with respect to either gene activation or repression

Histone marks typically function co-operatively with one another to either promote or repress the expression of genes. Acetylated histones often co-localise with H3K4me3 at the TSS and promoter region that results in a relaxed chromatin environment conducive the gene expression, while H3K27me3 and H2A.Z mono ubiquitination function co-operatively during polycomb mediated gene silencing in the body of repressed genes. To investigate these functional co-occurrences in our dataset we produced high quality, replicable, peak lists for each chromatin type. Differing strategies were used for narrow peaks and broad peaks, according to gold standard practise defined by the ENCODE consortium [46]. Irreproducible discovery rate (IDR) was used to call narrow peaks, which generated 19,704 (H3K4me3) and 13,590 (H3K56Kac) peaks for glandular trichomes, 18,668 (H3K4me3) and 14,147 (H3K56ac) for stem (internode), and 15,135 (H3K4me3) and 1,655 (H3K56ac) peaks for leaf tissue (Fig. 2a-b, Additional File 1: Fig. S7). Broad domain peaks were called by using reciprocal overlap between replicates, yielding 10,284 and 13,058 peaks for H3K27me3 and H2A.Z in *C. sativa* glandular trichomes respectively (Fig. 2c-d). All histone peaks for all tissue types were then annotated with respect to genomic location and gene structure (Additional Files 2-4).

**Fig. 2.**
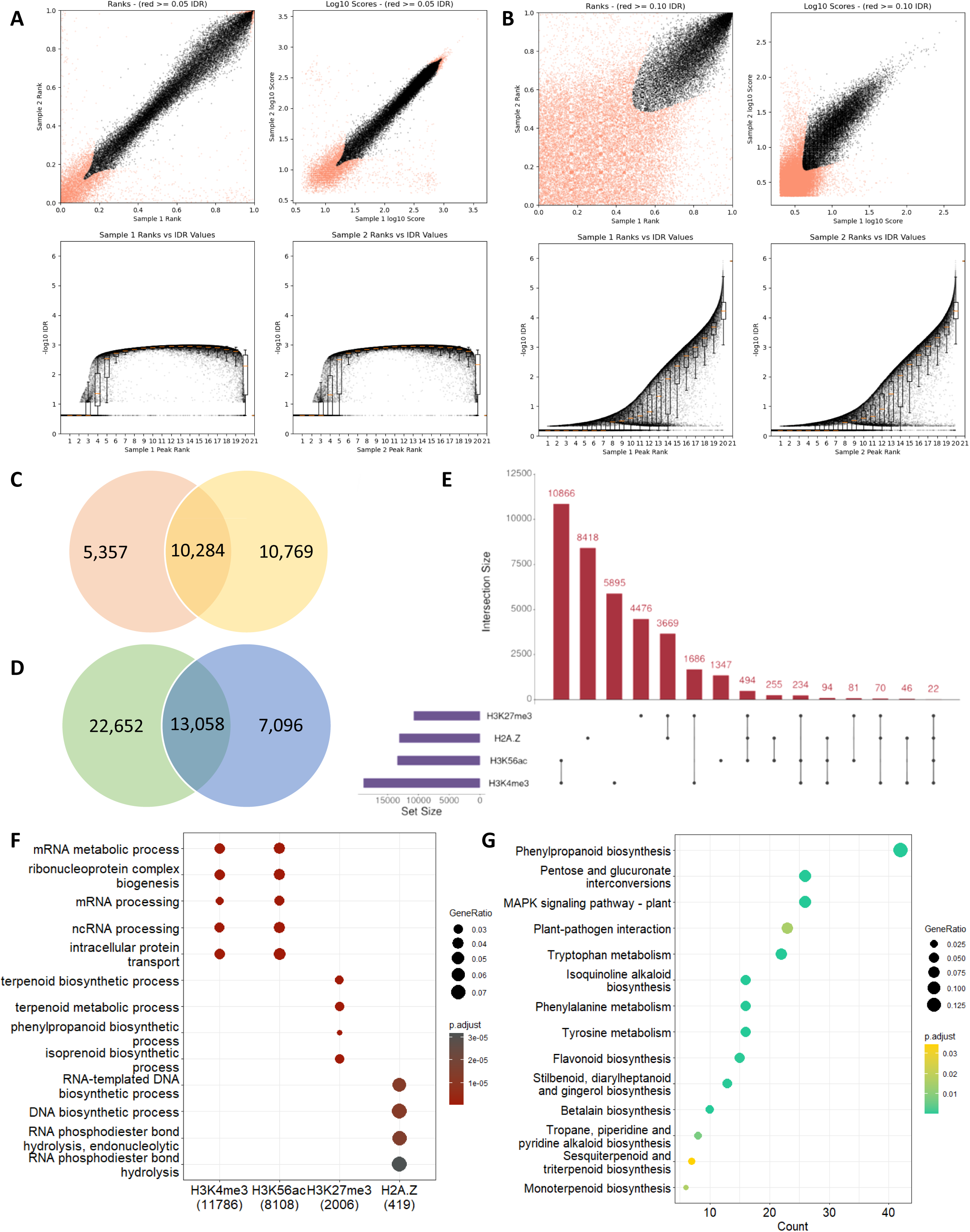
IDR analysis of **A** H3K4me3 and **B** H3K56ac yielded 19,704 and 13,590 replicable peaks respectively in glandular trichomes. BEDTools intersect analysis of **C** H3K27me3 and **D** H2A.Z yielded 10,284 and 13,058 replicable domains respectively in glandular trichomes. **E** Upstart plot shows histone modification co-localisation events in the glandular trichome genome. **F** Biological processes of genes containing H3K4me3, H3K56ac, H3K27me3, and H2A.Z in glandular trichomes. **G** KEGG ontologies of genes containing co-localised H3K4me3-H3K27me3 in glandular trichomes.

We investigated the potential co-occurrence in the genome of histone post-translational modifications within glandular trichomes (Fig. 2e). The majority of H3K4me3 and H3K56ac peaks co-occurred (10,866 overlapping peaks, compared to 5,895 individual H3K4me3 and 1,347 individual H3K56ac peaks), more than can be expected by chance (Fisher exact test p < 1×10^-5^). The second largest co-occurrence was between H3K27me3 and the histone variant H2A.Z (3,669 shared sites) (Fisher exact test p < 1×10^-5^), and substantial co-occurrence was also observed between H3K4me3 and H3K27me3 (1,686 overlapping peaks) (Fisher exact test p < 1×10^-5^) (Additional File 1: Figure S7b-d)

### The *C. sativa* glandular trichome epigenome indicates roles in plant specialised metabolism

Next, we asked whether different histone marks were associated with genes encoding specific functions, in glandular trichomes. Gene ontology (GO) enrichment analysis of biological processes (BP) was performed on genes found in each of the four chromatin types in glandular trichomes (Fig. 2f). There were 11,786 genes within H3K4me3 peaks and 8,108 within H3K56ac peaks. The genes associated with both histone marks were primarily enriched for housekeeping ontological terms including mRNA metabolic process, and mRNA processing. H3K27me3 and H2A.Z chromatin contained fewer genes (2,006 and 419, respectively). Genes found in H3K27me3 chromatin were enriched for GO terms related to plant specialised metabolism, such as terpenoid metabolic process and isoprenoid biosynthetic process in glandular trichomes. Genes associated with H2A.Z chromatin were enriched for GO terms associated with DNA homoeostasis and transcriptional control, in line with previous observations made in plants and animals (Additional File 5) [47, 48].

Bi-valent H3K27me3/H3K4me3 loci are involved in regulation of biosynthesis of the specialised metabolite camalexin in *A. thaliana* [17]. Our observations that H3K27me3 was frequently co-located with H3K4me3 in glandular trichomes, and that specialised metabolism genes were enriched within H3K27me3 regions, suggested that bi-valent chromatin states may be involved in regulation of specialised metabolism in glandular trichomes. We examined this by using the Kyoto Encyclopedia for Genes and Genomes (KEGG) curated pathways for *C. sativa* cs10 (GCF_900626175.2) to analyse pathways enriched amongst genes located in regions of overlapping H3K27me3/H3K4me3 chromatin. Pathways enriched amongst these genes were associated with plant specialised metabolism and defence, including phenylpropanoid biosynthesis, MAPK signalling, plant-pathogen interaction, flavonoid biosynthesis, and sesquiterpenoid biosynthesis (Fig. 2g, Additional File 6).

### *C. sativa* glandular trichomes display tissue specific epigenetic and transcriptional states linked to specialised metabolism

We reasoned that glandular trichome specific chromatin and transcripts could be identified using stem and leaf datasets to gain deeper insight into glandular trichome gene regulation. First, transcriptomes from leaf, stem and trichome were subjected to differential gene expression analysis in pairwise fashion for trichome vs. leaf and trichome vs. stem. In trichome vs leaf there were 1765 differentially expressed genes (log 2FC > 2, BH p.adj < 0.05), 1520 genes were significantly upregulated in trichomes while 245 were significantly upregulated in leaf. In trichome vs. stalk we found 1987 differentially expressed genes (log 2FC > 2, BH p.adj < 0.05), In trichomes there were 1448 upregulated genes and 540 upregulated genes in stem (Additional File 7). Next, we performed gene set enrichment analysis (GSE) of the differentially expressed genes (BP, molecular function, MF, and cellular compartment, CC). Genes differentially expressed in glandular trichomes, compared with both stem and leaf, were highly enriched for GO BPs including lipid metabolism, specialised metabolism, and organic acid biosynthesis in both cases (Fig.3a-b, Additional File 8). These genes were also enriched for GO MF terms including delta-9 tetrahydrocannabinolate synthase activity, cannabidiolate synthase activity, germacrene-a synthase activity, and other specialised metabolite related functions (Additional File 1: Fig. S8a-b, Additional File 8).

We then asked if glandular trichome specific differences in chromatin composition existed, if they contained genes, and whether different chromatin states were associated with genes of particular functions. To accomplish this, we derived the glandular trichome specific locations of each histone mark by comparison with the corresponding datasets generated for stem and leaf tissue, then curated gene lists where trichome specific peaks overlap genes in the cs10 genome as input for KEGG ontology analysis (Additional File 9). *C. sativa* glandular trichome specific H3K4me3 chromatin and H3K56ac chromatin was enriched in KEGG ontologies including starch and sucrose metabolism, sesquiterpenoid and triterpenoid biosynthesis, plant-pathogen interactions, and monoterpenoid biosynthesis (Fig. 3c-d). Glandular trichome specific H3K4me3-H3K27me3 bi-valent chromatin was enriched in KEGG ontologies including MAPK signalling, phenylpropanoid biosynthesis, and sesquiterpenoid and triterpenoid biosynthesis (Fig. 3e). Glandular trichome specific H3K27me3 chromatin and H2A.Z chromatin was enriched in KEGG ontologies related to biosynthesis of various plant specialised metabolites, phenylpropanoid biosynthesis, and mRNA surveillance (Fig. 3f-g). This analysis indicated that glandular trichomes associated processes including specialised metabolism are differentially regulated at the epigenetic level (Additional File 9).

**Fig. 3.**
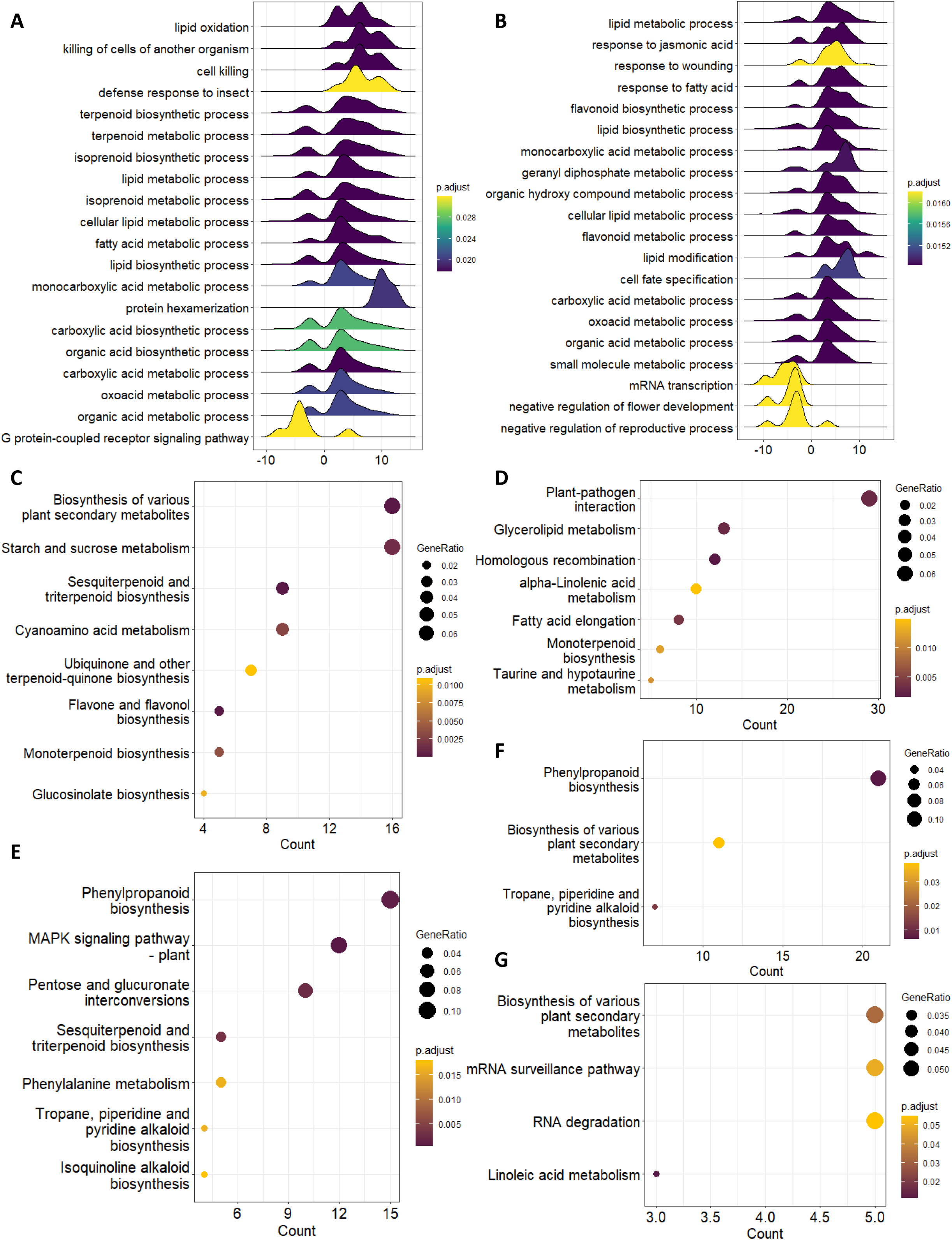
Tissue specific analysis of RNA and chromatin type datasets yield comprehensive insight into *Cannabis sativa* glandular trichomes. Pairwise gene set enrichment analysis (GSE) leveraging **A** Leaf and **B** Stem RNA datasets. KEGG ontology analysis of glandular trichome specific chromatin types **C** H3K4me3 **D** H3K56ac **E** H3K4me3-H3K27me3 **F** H3K27me3 and **G** H2A.Z.

We then asked if we could observe tissue specific differences in the chromatin composition of well characterised genes expressed specifically in glandular trichomes. For example, phytocannabinoid biosynthesis 2-acylphloroglucinol 4-prenyltransferase (also called cannabigerolic acid synthase, CBGAS/PT4; LOC115713185), ABC-transporter B family member 2 (LOC115716265), and alpha-humulene synthase (LOC115724563) were all associated with glandular trichome specific enrichment of the transcription-associated histone modifications H3K4me3 and H3K56ac in their promoter regions [9, 49]. The silencing marks H3K27me3 and H2A.Z were found in the gene body of alpha humulene synthase (LOC115724563) in stem and leaf tissues, consistent with the lack of expression of this gene in those tissues. H3K27me3 enrichment was less at alpha humulene synthase (LOC115724563) in glandular trichomes, but it was not completely absent. LOC115713185 and LOC115716265 were not associated with H3K27me3 or H2A.Z in stem or leaf, despite their lack of expression. Taken together, these observations may indicate further complexity or multivalency to chromatin states and the regulation of expression at individual genes in glandular trichomes (Additional File 1: Fig. S9a). The limitations imposed by analysing bulk samples of many cells cannot be fully excluded from our analysis but are mitigated against here by the analysis of highly pure glandular trichome samples.

### Glandular trichome specific epigenomic regulation of putative gene clusters in *Cannabis sativa*

Genes encoding components of specialized metabolite pathways are frequently found in plant gene clusters (GCs) [50]. These can be the consequence of local duplications such as gene family expansion in terpene synthases or, alternatively, unrelated genes that constitute or modulate a metabolic pathway may be co-located thereby forming a biosynthetic gene cluster (BGC) [51–56]. Genes in either cluster type are frequently expressed tissue or cell type specifically and corresponding to the tissues in which the product metabolite accumulates [57, 58]. This may indicate dynamic, long-range genome interactions in higher order topologically associating domains (TADS) that facilitate cell or tissue specific gene expression [59]. The epigenomic properties of such clusters are relatively poorly defined in plants and understanding them better may offer a unique and refined strategy to introduce entire biosynthetic gene pathways, discretely into a genome, in a tissue or cell type specific manner for applications in plant metabolic engineering.

We consequently examined the chromatin composition and expression status of genes located in putative *C. sativa* gene clusters, focusing upon two clusters that may have glandular trichome specific activity. We drew upon our previous computational predictions of BGCs in this *C. sativa* [60]. Putative BGC #17 (monoterpene biosynthesis) exhibited glandular trichome specific gene expression, and enrichment of transcriptionally active chromatin (H3K4me3 and H3K56ac), in limonene synthase/TPS1 (LOC115716064), and two myrcene synthases (LOC115716063, LOC115716405) (Fig. 4a). Interestingly, enrichment of H3K27me3 at these genes in leaf and stem tissue, where they are not expressed, implies a role for polycomb mediated gene silencing in putative BGC #17. Contrastingly, putative BGC #28 had trichome specific gene expression of class V chitinases *CHIT5* and *CHIT5*-like (LOC115724705 and LOC115695573), germacrene-A synthase (LOC115695573), and ferredoxin-NADP reductase embryo like (LOC115724444) but did not exhibit clear relationships between histone modifications H3K4me3 and H3K56ac and transcription. The chromatin state at germacrene-A synthase (LOC115695573) was indistinguishable between either tissue at the epigenomic level, except for more pronounced H2A.Z deposition near the promoter region. Ferredoxin-NADP reductase embryo-like (LOC115724444) was expressed solely in the glandular trichomes, yet there was no discernible difference in either H3K4me3 or H3K56ac deposition between glandular trichomes and other tissues. There was however a subtle enrichment of H2A.Z in the gene body of LOC115724444 in glandular trichomes. Furthermore, throughout the length of BGC #28 there was more pronounced enrichment of H2A.Z in trichomes comparatively, a feature which has been associated with tissue specific BGC regulation in *A. thaliana* (Additional File 1: Fig. S9b) [61].

**Fig. 4.**
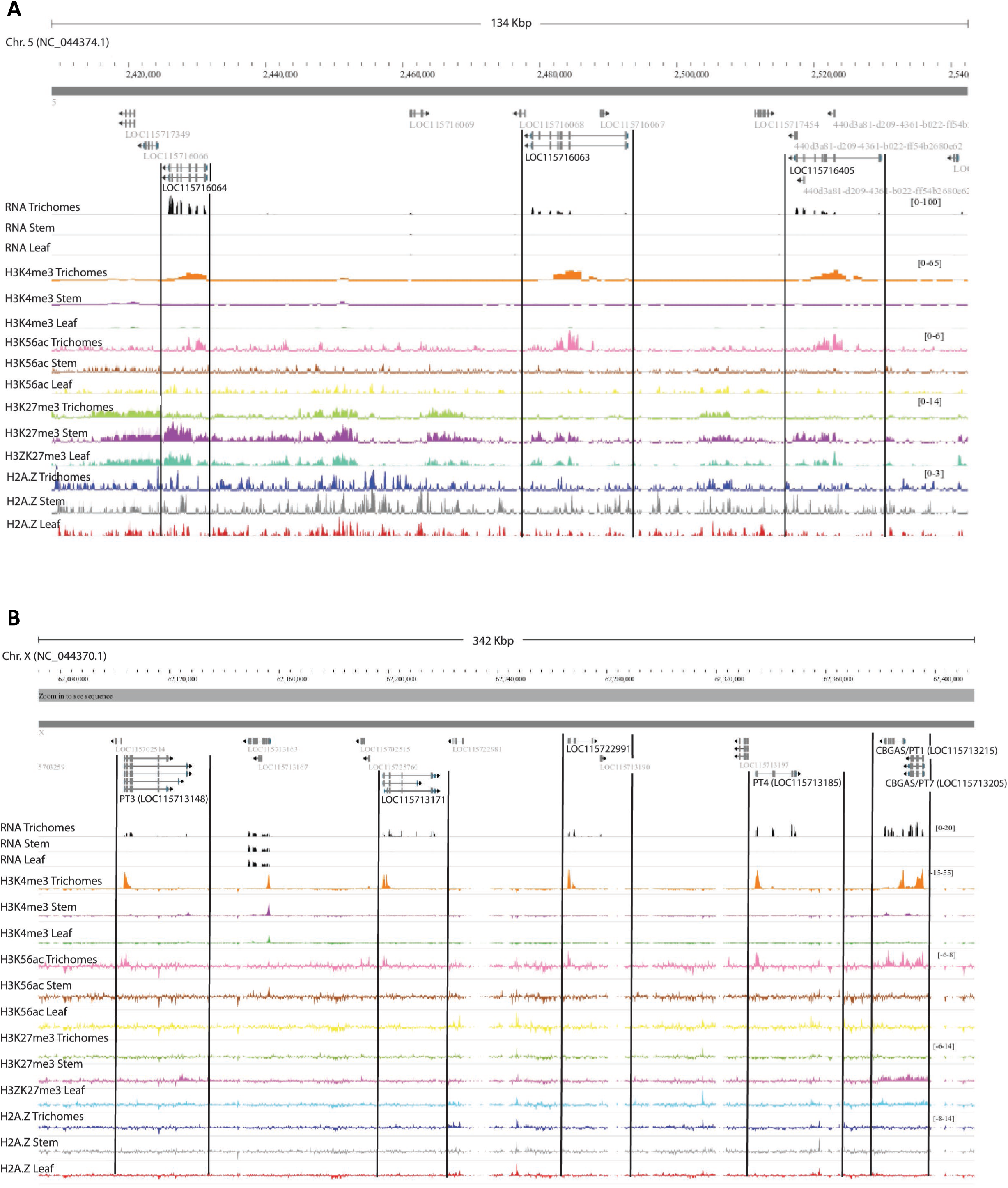
**A** Monoterpene synthase gene cluster previously identified in Conneely et al. 2022 as CBDRx cluster no.17 shows glandular trichome specific gene expression and chromatin landscape. **B** Genome data viewer screenshot of the 342 Kbp region encompassing the cannabigerolic acid synthase / prenyltransferase gene cluster in C. sativa. The CBGAS cluster genes PT3 (LOC115713148), LOC115713171, LOC115722991, PT4(LOC115713185), CBGAS/PT1(LOC115713215), CBGAS/PT7(LOC115713205) show trichome specific expression and enrichment of H3K4me3 and H3K56ac at their promoter regions.

In our data we also saw evidence of trichome specific epigenomic and transcriptomic regulation of a novel putative cluster of six aromatic prenyltransferases (Fig. 4b, Additional File 7). This included the cannabigerolic acid synthases PT1, PT4, and PT7 (LOC115713215, LOC114713185, and LOC115713205) cannaflavin biosynthesis gene PT3 (LOC115713148) as well as two additional uncharacterised CBGAS-like genes (LOC115713171 and LOC 115722991) in a 347 Kbp region of the X chromosome [62, 63]. This cluster was not detected in our analysis of BGCs however, considering the homology and neo-functionalisation, this cluster is likely the result of ancestral gene expansion, a homologous gene cluster [60]. These genes had glandular trichome specific enrichment of H3K4me3 and H3K56ac in their promoters compared to other intervening genes, consistent with their active transcription in that tissue.

### Identification of putative glandular trichome specific regulatory motifs using H3K56ac enrichment

Active plant cis-regulatory elements (CREs) are often flanked by H3K56 acetylated chromatin [30]. CREs typically contain TF binding sites that drive target gene expression through interactions with target gene promoters, either because the binding site is within the promoter itself or through long-range promoter-enhancer interactions. We reasoned we could use H3K56ac data for trichomes, stem, and leaf to mine CREs with glandular trichome specific activity. To this end, we examined H3K56ac chromatin loci specific to glandular trichomes. We focused on identifying novel gene distal and intragenic CREs that cannot be located by simple motif enrichment analysis of gene promoter regions. This was achieved by excluding peaks proximal to gene promoters (≤1.5 Kbp from TSS). We identified 1,048 H3K56ac loci representing putative gene distal and intragenic CREs specific to glandular trichomes. Putative CREs were significantly enriched for 65 DNA sequence motifs (q < 0.05, JASPAR CORE database). The enriched motifs included MYB, MYB-related, NAC, and TCP transcription factor binding motifs (Fig. 5a, Additional File 10).

**Fig. 5.**
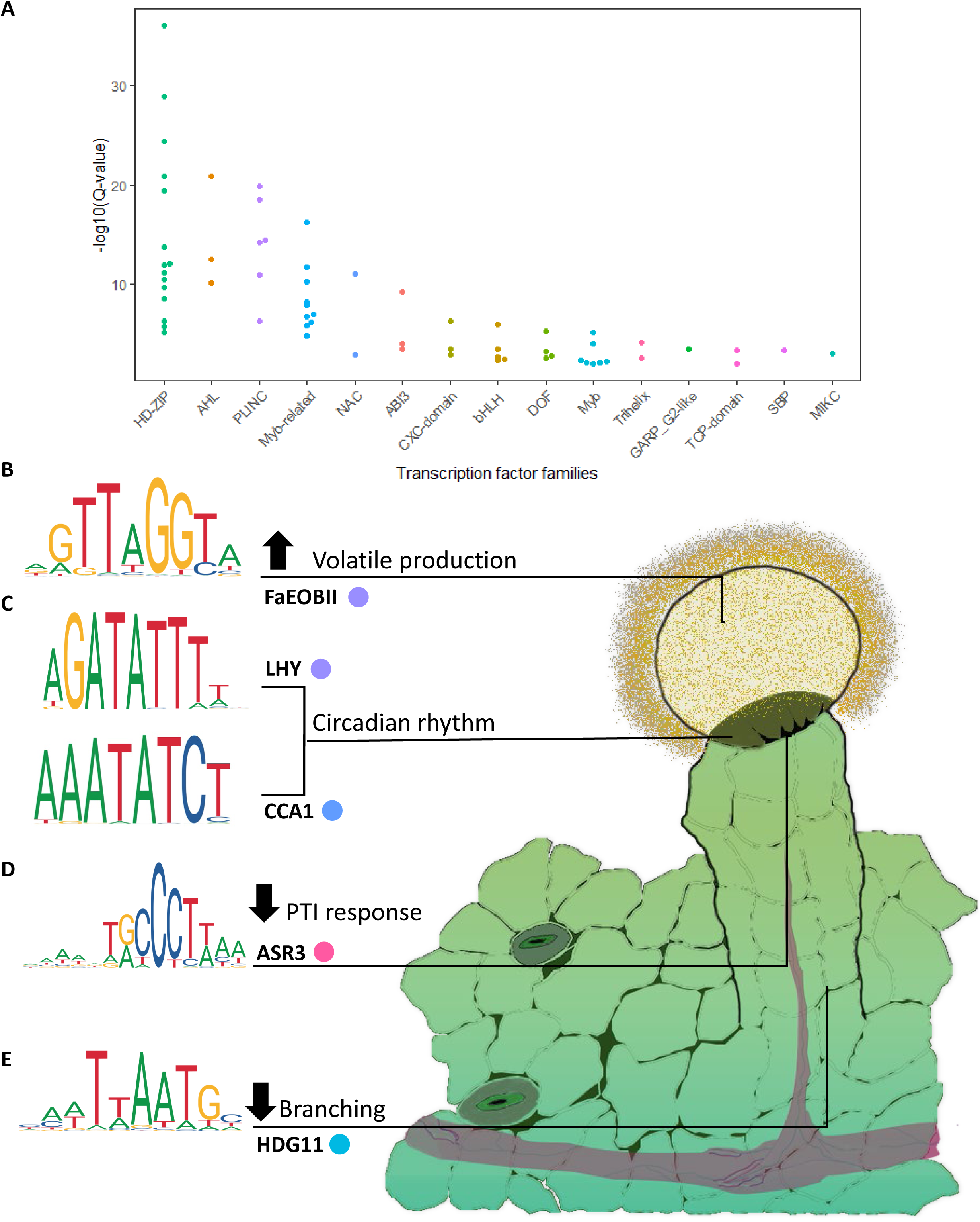
**A** Beehive plot showing various transcription factor families associated with mined motifs in glandular trichomes specific cis-elements. **B** FaEOBII motif enriched in glandular trichome cis-element candidates; schematic highlights known role in aroma volatile production **C** LHY and CCA1 motifs; circadian rhythm is known to be linked to cannabinoid production **D** ASR3 motif is associated with pattern triggered immunity in plants **E** HDG11 motif and its cognate transcription factor are known to reduce trichome branching consistent with the lack of branching observed along the multicellular stalk.

We examined the cognate transcription factors that may recognise these motifs and examined their likely functions, to better understand how they may be related to glandular trichome biology. For example, the MYB-related FaEOBII cis element was enriched in putative CREs (Additional File 10). The corresponding best blast P hit for its associated transcription factor in the cs10 reference assembly was the MYB-related transcription factor CsEOBI (LOC115699015). CsEOBI is highly expressed in the glandular trichomes compared to either leaf (log2FC = 7.49, BH p.adj = 0.000105) or stem (log2FC = 8.08, BH p.adj = 1.21E-05). FaEOBII is known to positively regulate production of volatile organic compounds (Fig. 5b) [64].

Interestingly, LHY and CCA1 motifs were also enriched in putative CREs. The cognate transcription factors for these motifs work cooperatively to regulate circadian rhythms (Fig. 5c) [65]. The trihelix family member ASR3 motif was enriched, and its associated transcription factor negatively regulates pattern triggered immunity (PTI) that mitigates unnecessary hypersensitivity responses (Fig. 5d). HDG11 motifs were also enriched in glandular trichome specific H3K56ac chromatin. HDG11 has been shown to negatively regulate trichome branching in other plant species (Additional File 10).

### Predicting individual enhancer-gene interactions specific to glandular trichomes

We reasoned that a population of tissue specific enhancer sequences, which drive gene expression in glandular trichomes, would be amongst the identified 1,048 gene distal and intragenic CREs. To try and identify these we used a nearest gene model by matching the nearest gene (TSS) to each 1,048 predicted CREs. We then refined these gene-enhancer candidates including only those with genes expressed in the glandular trichomes specifically. This identified 87 putative CREs whereby the closest neighbouring gene in cis showed considerably greater expression in glandular trichomes relative to both stem and leaf tissue (log2FC >2, BH p.adj<0.05) (Table 1, Additional File 11). Strikingly, this strategy captured gene distal and intragenic H3K56ac associated with the expression of hallmark glandular trichome associated genes. These included genes involved in terpenoid specialised metabolism, such as sesquiterpene biosynthesis; alpha-humulene synthase (LOC115724563), monoterpene biosynthesis; myrcene synthase (LOC115716405) and limonene synthase (LOC115716064), triterpene biosynthesis; beta-amyrin synthase (LOC115719748). They also included genes involved in cannabinoid specialized metabolism, such as biosynthesis of the cannabinoid precursor hexanoate Butanoate-CoA ligase AAE1 (LOC115713865), as well as cannabidiolic acid synthase (LOC115697762). Fatty acid biosynthetic genes implicated in the upstream production of fatty alkyl side groups required for cannabinoid production were also found (Lineolate 13S-lipoxygenase, LOX1, LOC115719612; Delta (12)-oleate desaturase, FAD1, LOC115719329), as were specialised metabolite transporters (ABC transporter B family member 2, LOC115716265; ABC transporter G family member 20, LOC115711415), and a transcription factor involved in trichome maturation (MYB106, LOC115701410). Interestingly, several trichome specific gene-enhancer predictions identified genes involved biotic stress resistance that have not yet been reported in studies of glandular trichomes nor *C. sativa* including disease resistance protein RMP1 (LOC115704534) that is currently being investigated in wheat for its properties in resistance to powdery mildew, and an unusual protease neprosin (LOC115720846) found in carnivorous plants.

**Table 1.**
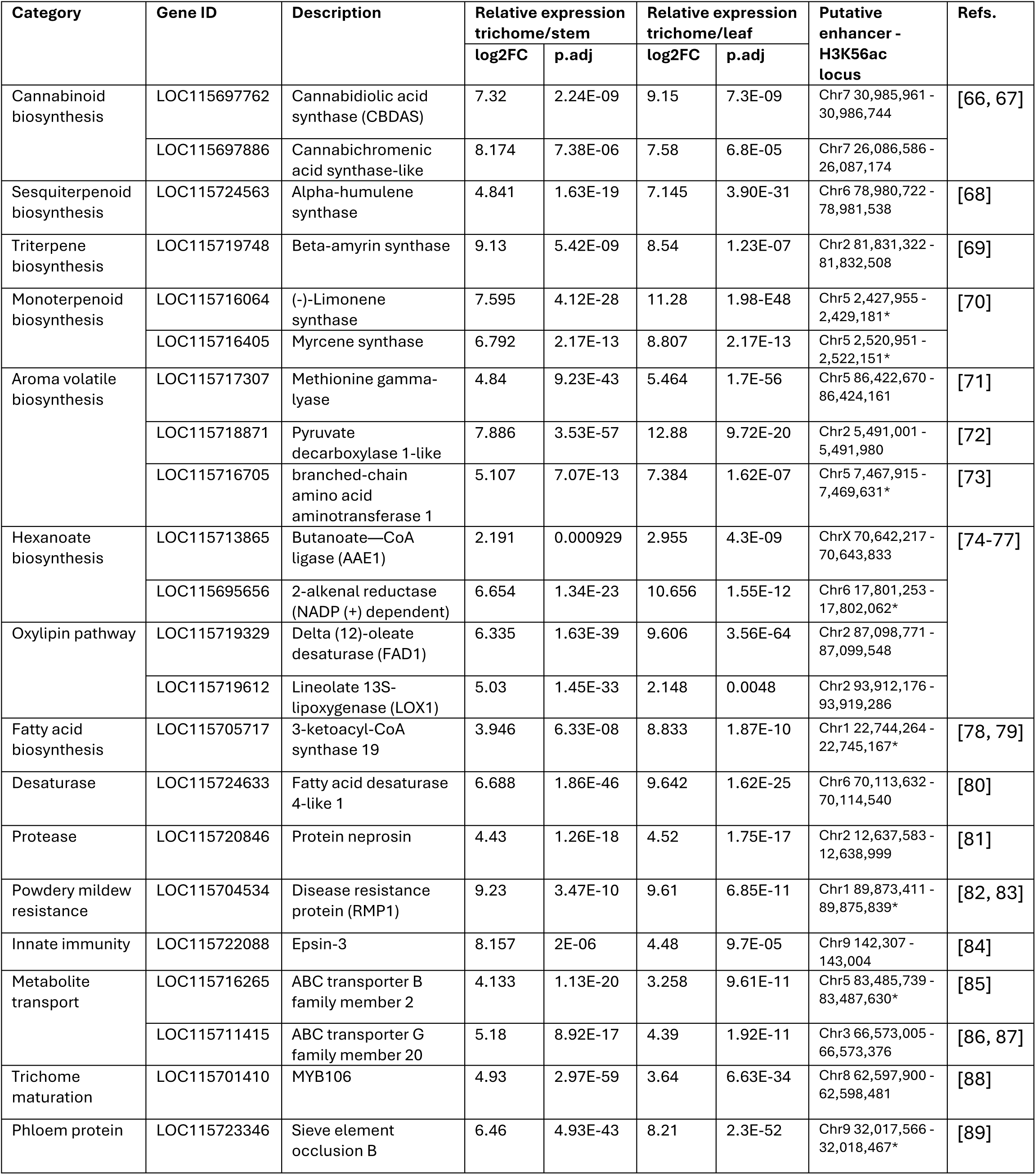
Closest neighbouring genes that are expressed glandular trichome specifically to *C. sativa* glandular trichome specific gene distal H3K56ac loci, showing expression – a list of putative tissue specific enhancer gene interactions. * Indicates intragenic putative cis-regulatory elements.

## Discussion

*Cannabis sativa* is a plant with significant biotechnological potential owing to its specialised metabolite productivity and large capitate stalked glandular trichomes. Despite our reliance on plant specialised metabolite products in industry, food, and medicine, our understanding of glandular trichomes is disproportionately sparse. Currently, only a single publication in plants investigates epigenomic regulation of plant glandular trichome activity [90]. Here, we have provided the first H3K4me3, H3K56ac, H3K27me3, and H2A.Z chromatin maps in *C. sativa* glandular trichomes, and in plant glandular trichomes more broadly. Additionally, we demonstrate that classical eukaryotic functions of histone post translational modifications are functionally conserved in the *C. sativa* epigenome (Fig. 1).

Our chromatin landscape maps have been assembled using gold standard ENCODE guidelines to provide confidence in the observations made in this study, as well as to enable further research by the plant science community wherein such datasets are not publicly available. Datasets such as these provide a richer understanding of the biological intricacies that govern gene expression and chromatin dynamics *in planta*. A key theme we observe in this study is that chromatin modifications often co-localise, presumably to co-operatively regulate gene expression. For example, H3K56 acetylated chromatin and H3K4me3 co-localize at the promoters and TSSs of actively transcribed genes, whilst H3K27me3 and H2A.Z co-localize in the gene bodies of silenced genes (Fig. 2).

Our analysis highlights tissue-specific gene expression and chromatin states consistent with focal points of specialised metabolism, carbon trafficking, and plant abiotic and biotic stress resistance, as would be expected given the functions of glandular trichomes. For example, we observed that glandular trichome specific putative H3K4me3-H3K27me3 bi-valent chromatin is enriched in specialised metabolism, defence signalling, and innate immunity associated genes. We speculate that complex chromatin landscapes like these have the potential to integrate environmental cues to dynamically allocate carbon resources as needed during defence response elicitation like that reported in camalexin biosynthesis (Fig. 3) [17]. It should be noted, however, that overlaps between H3K4me3-H3K27me3 loci in leaf or stem tissue might reflect the heterogenous composition of those tissues, as opposed to a bona-fide signal of bi-valency. This limitation is less probable in glandular trichomes samples, which were isolated to a high degree of purity largely composed of disc cells.

Chromatin landscapes regulate plant BGC expression, so examining these landscapes provides an opportunity to better define *C. sativa* BGCs and understand how their activity may be regulated [91, 92]. We investigated our previously predicted putative *C. sativa* BGCs for evidence of trichome specific gene expression and chromatin composition [60]. We observed that two putative BGCs (#17, terpene-related; #28, terpene and fungal resistance related) from our previous study were expressed specifically in glandular trichomes. Most genes in both BGCs displayed enrichment of promoter H3K4me3 and H3K56ac specifically in glandular trichomes, consistent with active gene expression. Observations made in *A. thaliana* suggest enrichment of H2A.Z is functionally associated with active BGCs in tissues where they are expressed, and enrichment of H3K27me3 at those BGCs in tissues where they are not expressed [93]. Contrasting with this, we did not observe enrichment of H2A.Z in a tissue specific manner that might explain trichome specific expression of either BGC. However, we did observe enrichment of H3K27me3 in BGC #17 monoterpene genes in stem and leaf tissue, where these genes were not expressed. Nonetheless, our data prompts further research into the association of distinct H2A.Z and H3K27me3 chromatin states at active and repressed BGCs. Future investigation might also consider the capacity of H2A.Z, a histone variant, to be post-translationally modified itself [45].

We made use of the association between H3K56ac and CREs to identify putative *C. sativa* glandular trichome specific TFs and binding motifs [30, 31, 94]. This strategy yielded candidates involved in aroma volatile production associated (FaEOBII, JASPAR ID MA1408.1), trichome development (HDG11; MA0990.1), and circadian rhythms (LHY; MA1185.1, CCA1; MA0972.1). Daytime progression has recently been implicated in cannabinoid production which might explain the enrichment of circadian rhythm associated motifs in our dataset (Fig. 5) [95]. Furthermore, we hypothesised that we could leverage our H3K56ac and gene expression datasets to mine the glandular trichome epigenome for gene distal and intragenic (≥1.5 Kbp from TSS) loci that likely contain enhancers influencing the closest gene. We note that enhancer elements may also regulate genes at very long range, acting over intervening genes, and so this approach would only yield a subset of the potential CREs in the genome. Nonetheless, our approach yielded a large repertoire of enhancer-gene pairs where the genes have known glandular trichome specific functions, including CBDAS (LOC115697762), terpenoid synthases (LOC115724563, LOC115719748, LOC115716064, LOC115716405), and metabolite transporters (LOC115716265, LOC115711415) [66, 87, 96, 97]. Moreover, the data brought highlighted a glandular trichome specific alkenal NADP reductase (LOC115695656) that is a candidate for the twice postulated alkenal NADP reductase involved in the biosynthesis of hexanoate and production of cannabinoids downstream [76, 77]. Our approach was limited to 2-dimensional space, and therefore could not consider potential 3D interactions between enhancers and cognate promoters in cis and trans, that would likely identify potential interactions for many more of the putative CREs we predicted. Further study of these would require assays like Hi-C to capture the full scope of enhancer gene interactions in *C. sativa* glandular trichomes. Nonetheless, the putative enhancer-gene interactions we identify here in *C. sativa* presents a significant biotechnological resource that may provide a framework for engineering glandular trichome gene expression in the future.

## Conclusion

In this study we provide novel insights into the chromatin landscape of glandular trichomes through the analysis of multiomic datasets. Our work shows how chromatin features and function are conserved in *C. sativa* compared with more deeply characterised plant species, from feature types to correlation with gene expression, deposition and localisation within genes, and association with distal regulatory regions. Moreover, we show the advantages of studying such chromatin features where, for example, genes postulated to exist in previous studies have been predicted by our multimodal approach, and genome wide sequence information can be mined in tissue specific manners to identify cis-regulatory elements not yet observed in *C. sativa*. This work has laid a solid foundation for our understanding of glandular trichome chromatin dynamics with respect to tissue specific gene expression and cis-regulatory elements, whereby the product of studies such as this will be at the core of glandular trichome metabolic engineering in years to come.

## Methods

### Plant Material and Growth Conditions

Cuttings were taken from female genotype (MW6-15) industrial hemp line (Accession #6) mother plants. Cuttings were made using a sterile blade and at a 45-degree angle immediately below the 3^rd^ node from the apical meristem. The cuttings were partially pruned – removing leaf tips and large leaves, taking care not to remove all leaves. The cuttings were then dipped in 3.0 g/L indole butyric acid rooting gel hormone (Growth Technology © Clonex purple) and allowed to sit for 30 seconds. Cuttings were individually transferred to Grodan Rockwool cubes (36mm x 36mm x 40mm) previously saturated with half strength CANNA veg fertiliser (20 mL A + 20 mL B/ 10 L water), and then transferred to propagator humidity domes (4 clones to one propagator) with the vent fully closed. Clones were monitored daily – half strength CANNA veg fertiliser (20 mL A + 20 mL B/ 10 L water) was added to the base of the propagator when required and vents were opened in small daily increments until completely opened and grown under vegetative lighting conditions of 18 hours on 6 hours off using Philips Master TL-D Super 80 low-pressure mercury discharge lamps. Clones were kept in propagators until established roots could be observed (approximately 2 weeks) at the base of the rockwool cubes. Potting mix containing 1:1:1 peat moss, perlite, and vermiculite supplemented with 1 g/L dolomite was prepared and divided into 1 L pots. Pots were then saturated with full strength CANNA veg fertiliser (40 mL A + 40 mL B/ 10 L water). Clones were then transferred to appropriately labelled 1 L pots and allowed to grow under vegetative lighting conditions for 4 weeks. Plants were monitored daily and watered regularly using full strength CANNA veg during the vegetative growth stage.

Following 4 weeks of vegetative growth plants were re-potted into appropriately labelled 10 L pots using the above potting mix recipe. Potting mix was saturated with full strength CANNA flora (40 mL A + 40 mL B/ 10 L water) prior to re-potting of the clones. The lighting cycle was then varied to 12 hours on and twelve hours off to induce flowering. Clones were grown under flowering conditions for 6 weeks and watered daily with full strength CANNA flora. Clones were harvested during the late stage of flowering as defined by a colour change, from white to brown, of greater than two thirds of all stigmas on the plant.

### Tissue harvest and trichome isolation for RNA sequencing

For RNA extraction, fresh samples were collected in three biological replicates from mature leaves, stem (internode), and female inflorescences. Triplicates were obtained from three individual plants. Trichomes were isolated from female inflorescences using a method modified from a protocol previously described [98]. Briefly, ∼5 g samples were transferred to 50 mL Falcon^TM^ tubes and about 10 mL of liquid nitrogen was added to each tube. The tubers were loosely capped and vortexed until the trichomes were fully removed. After removing plant debris by inverting and tapping the tubes, trichome-enriched samples were carefully transferred to 2 mL Eppendorf tubers in liquid nitrogen for further processing.

### Tissue harvest and trichome isolation for chromatin immunoprecipitation

Samples of stem (internode) and vegetative leaves were taken from each clone using a sterile blade. Each sample was transferred to an appropriately labelled 50 mL falcon tube, sealed, and immediately flash frozen in a Dewar containing liquid nitrogen. Samples were then stored at - 80°C.

Glandular trichomes were isolated by ice-water extraction – all inflorescences were harvested from an individual clone and cut into small 5 cm x 5 cm pieces using sterile secateurs. Nylon bags with 25 µm, 45 µm, 73 µm, 120 µm, and 160 µm mesh sizes were sequentially placed inside one another starting the smallest, outermost, 25 µm mesh bag until the largest mesh size, innermost, 160 µm mesh bag. The setup was then placed in a 5 L beaker and then filled through the innermost nylon mesh bag with 1:3 ice-cold milli-q water, crushed ice. The cut inflorescences were then added to the beaker/nylon-mesh set-up and stirred using a large metal spatula for ten minutes. The nylon-mesh bags were then removed from the beaker and the contents of the beaker were then filtered through a nylon mesh with 25 µm pore size.

Glandular trichomes were then retrieved, *via* pipetting, from the surface of the 25 µm nylon mesh and transferred into appropriately labelled falcon tubes. Glandular trichomes were immediately frozen in liquid nitrogen and then transferred to cold storage at −80°C.

### Tissue preparation and chromatin cross-linking

Glandular trichomes, stem (internode), and vegetative leaves were retrieved from cold storage (−80°C) and approximately 1 g of each tissue was ground into a fine powder using liquid nitrogen and a pre-chilled mortar and pestle. The ground tissues were then transferred into appropriately labelled 15 mL falcon tubes followed by the addition of 12.5 mL of nuclear isolation/ cross-linking buffer (60 mM HEPES at pH 8.0, 1 M Sucrose, 5 mM KCl, 5 mM EDTA at pH 8.0, 0.6% (v/v) Triton X-100, 1 mM PMSF (Sigma-Aldrich 93482-50ML-F), 1 mM pepstatin A (Sigma-Aldrich P5318-5MG), 1 mini-complete tablet (Sigma-Aldrich 11836170001) per 10 mL buffer) on ice (Additional File 12: Table A1). The contents of each tube were stirred until a homogenous suspension had formed. Next, 360 µL of 37% formaldehyde was added to each of the sample suspensions - cross-linking was achieved through incubation for 25 mins at room temperature and gentle rotation. Cross-linking was halted through the addition of 875 µL 2 M glycine to each of the sample suspensions followed by 25 mins incubation at room temperature and gentle rotation.

### Nuclei Isolation

Nuclei were isolated from the sample suspensions through passive filtration into a 50 mL falcon tube using a 40 µm nylon mesh sieve. The filtrate was then centrifuged at 4000 RPM for 20 mins at 4°C. The supernatants were carefully removed, and the soft nuclei pellets were resuspended, by pipetting, using 1 mL of extraction buffer (0.25 M Sucrose, 10 mM Tris-HCl at pH 8.0, 10 mM MgCl2, 1%(v/v) Triton X-100, 1 mM EDTA at pH 8.0, 5 mM β-mercaptoethanol, 1 mM PMSF, 1 mM Pepstatin A, 1 mini-complete table per 10mL)( Additional File 12: Table A2) and then transferred to a 2 mL Eppendorf tube. The walls of each 50 mL falcon tube were washed twice with 100 µL of extraction buffer and transferred to its corresponding Eppendorf tube to collect any residual nuclei. Tubes were then sealed and centrifuged at 11.4k RPM for 10 mins at 4°C and the supernatant discarded.

### Chromatin Shearing

Each of the pellets were then resuspended in 300 µL of nuclei lysis buffer (50 mM Tris-HCl at pH 8.0, 10 mM EDTA at pH 8.0, 1% (w/v) SDS, 1 mM PMSF, 1 mM Pepstatin A, 1 mini-complete table per 10 mL buffer) (Additional File 12: Table A3). Samples were then sonicated using a Diagenode Bioruptor® on high setting, 30 seconds on, 30 seconds off, for 15 mins at 4°C. Samples were then centrifuged at 5000 RPM for 10 minutes at 4°C to pellet unwanted cellular/nuclear debris. The supernatant, containing sheared chromatin, was then transferred to new Eppendorf tubes on ice. All samples were then placed in cold storage at −80°C.

### Chromatin Immunoprecipitation

In preparation for immunoprecipitation 60 µL of Dynabeads-Protein A (Thermo Fisher Scientific, #10002D) per sample per histone mark was washed with 30 µL of ChIP dilution buffer (1.1% (v/v) Triton X-100, 1.2 mM EDTA at pH 8.0, 16.7 mM Tris-HCl at pH 8.0, 167 mM NaCl, 1 mM PMSF, 1 mM Pepstatin A, 1 mini-complete tablet per 10 mL buffer) (Additional File 12: Table A4). Samples were pre-cleared by adding 90 µL of the washed Dynabeads-Protein A and incubating for 4.5 hours, rotating, at 4°C. Eppendorf tubes were labelled with the appropriate sample name and histone mark and 60 µL aliquots of the washed Dynabeads-Protein A were made into each tube followed by the addition of 2.5 µL (2.5 µg) of either Anti-H3K4me3 (Millipore®, 07-473), Anti-H3K56ac (Millipore®, 07-677-1), Anti-H3K27me3 (Millipore®, 07-449), or Anti-H2A.Z (Millipore®, 07-594) to their correspondingly labelled tubes. The tubes were then sealed and incubated at 4°C, rotated, for 1 hour.

Following pre-clearing, samples were then placed on a magnetic stand (Thermo Fisher Scientific, #AM10027) for 2 mins until all Dynabead-Protein A precipitate out of solution. Next, 300 µL of the supernatant, of each tissue type, was then transferred to a new tube to use as an input control. One input control was made for each of the tissue types used in the experiment. The INPUT controls were then immediately placed in storage at −20°C. The remaining 3.6 mL of supernatant was then made into 4 x 900 µL aliquots and transferred to the previously labelled, tubes, containing the Dynabeads-Protein A/antibody slurry and incubated at 4°C, rotating, for 90 mins. The samples were then placed on a magnetic stand for 2 mins and the supernatant was removed. On the stand, 1 mL of low salt wash buffer (150 mM NaCl, 0.1% (w/v) SDS, 1% (w/v) TritonX-100, 2 mM EDTA at pH 8.0, 20 mM Tris-HCl at pH 8.0) was added to each tube (Additional File 12: Table A5). The tubes were then sealed, and the contents were resuspended by inverting and then placed on the magnetic stand for 2 mins until all the Dynabeads-Protein A/antibody/chromatin complexes precipitate out of solution and the supernatant discarded.

The wash step was repeated one more time using the low salt wash buffer, then again using the high salt wash buffer (500 mM NaCl, 0.1% SDS, 1% TritonX-100, 2 mM EDTA at pH 8.0, 20 mM Tris-HCl at pH 8.0) (Additional File 12: Table A6), followed by a final wash using the LiCl wash buffer (0.25 M LiCl, 1% (v/v) NP-40 (Sigma-Aldrich NP40S-100 mL), 1% Sodium Deoxycholate, 1 mM EDTA at pH 8.0, 10 mM Tris-HCl at pH 8.0) (Additional File 12: Table A7). The beads were then washed using 1 mL of TE buffer (10 mM Tris-HCl at pH 8.0, 1 mM EDTA at pH 8.0) (Additional File 12: Table A8), allowed to precipitate on the magnetic stand for 2 mins, followed by aspiration of the supernatant. Each sample was then removed from the magnetic stand and the pellets were resuspended with 150 µL of SDS elution buffer (1% SDS, 0.1 M NaHCO_3_) (Additional File 12: Table A9) and incubated at 65°C for 15 minutes. The samples were then placed on the magnetic stand for 2 mins and the supernatant was then transferred to a newly labelled 1.5 mL tube.

A second elution of the beads was performed using 150 µL of SDS elution buffer. The supernatants were combined in 1.5 mL tube for a final volume of 300 µL. A master mix solution containing 12 µL 5M NaCl, 30 µL Dithiothreitol (DTT), and 30 µL 1M NaHCO3 per sample was made up for all samples, including INPUT controls. 72 µL of the master mix was aliquoted into each of the samples and INPUT controls and allowed to incubate overnight at 65°C. Samples and INPUT controls were retrieved from overnight incubation and 6 µL 0.5M EDTA, 12 µL 1M Tris pH 7.0, and 2 µL proteinase K was added to each tube. Tubes were then incubated at 45°C for 1 hour. Samples and INPUT controls were then cleaned up by adding 350 µL chloroform/ isoamyl alcohol (24:1), vortexing, and centrifuging using a bench top centrifuge at max speed for 25 mins at room temperature. The top layer was then carefully, without disturbing the interphase, transferred to a new tube. 2 µL of glycogen, 60 µL of 3M sodium acetate pH 5.2, and 900 µL of chilled ethanol (95%) was added to each sample and INPUT control and then allowed to precipitate overnight at −20°C. Samples and INPUT controls were then centrifuged at 16,000 g, for 30 mins, at 4°C. The DNA pellet was then washed two times using 1 mL of ethanol (70%). The washed DNA pellets were then allowed to dry in a fume hood. The dry DNA pellets were then resuspended in 50 µL TE buffer. Double stranded DNA (dsDNA) content was quantified with the Qubit dsDNA high sensitivity assay using the Qubit 4 fluorometer (See Additional File 1). ChIP-seq libraries were then generated using the Accel-NGS 2S Plus DNA library Kit (Swift biosciences), following the manufacture’s recommendations using 10 pg – 250 ng of input dsDNA per sample.

### Assessing library quality – Tapestation

The quality of the sequencing libraries was assayed using a 2200 Tapestation (Agilent) and D1000 screen tape. Samples were prepared using 1 µL of each sample library and diluted using 3 µL of D1000 sample buffer for a final volume of 4 µL. Tubes were then sealed and vortexed for 30 seconds. The samples were then spun down in a microfuge for 1 min to ensure residual droplets on the tube walls were collected at the base of each tube. Sample were then run on the Tapestation, and the results recorded.

### Chromatin immunoprecipitation sequencing operations

Samples were sequenced on an Illumina next-seq according to manufacturer’s instructions. We applied the ENCODE consortium guidelines for broad (H3K37me3, H3K56ac, H2A.Z) and narrow peak (H3K4me3) type data in humans of >45 million reads and >20 million reads are recommended respectively. For *C. sativa* the target reads were linearly scaled approximately 4 times to >10 million reads and >5 million reads respectively to account for the genome size discrepancy between human and *C. sativa*.

### RNA sequencing library preparation and sequencing operations

All samples were homogenised using Geno/Grinder 2010 (SPEX SamplePrep) and total RNA was isolated from homogenised samples using the Sigma Spectrum Plant Total RNA kit (Sigma) supplemented with the On-Column DNase I Digestion step to remove genomic DNA. RNA was eluted in 50 μL EB buffer. RNA concentration was measured using a Nanodrop spectrophotometer. RNA-seq library generation was performed using Illumina TruSeq Stranded mRNA Library Prep and indexed using Truseq RNA UD Indexes (96 indexes, 96 samples). Individual libraries were quality checked using Qubit dsRNA HS Assay kit and Agilent Tapestation (D1000) before being pooled into one sample for sequencing using Illumina NextSeq 500/550 High Output kit v2 (75 cycles).

### ChIP-seq data processing

Raw sequencing reads were quality trimmed using Trim Galore version 0.6.3 and the following parameters were used for paired end reads -paired -trim1 -fastqc [99]. The -fastqc option was selected to provide a quality report of the trimmed reads after trimming.

Quality trimmed reads were then aligned to the cs10 version 2 reference genome GCF_900626175.2 using Bowtie2 version 2.3.5.1 using the default parameters for paired end reads. Bowtie2 output SAM files were processed using Samtools version 1.9. SAM files were first converted to BAM files using the Samtools view -Sb function. Samtools view output BAM files were then sorted using the Samtools sort function [100]. The aligned reads were then filtered for PCR duplicates. Using the Picard version 2.2.2 suite of tools PCR duplicates were marked and subsequently removed from the aligned files using the MarkDuplicates function with the following parameters: TAG_DUPLICATE_SET_MEMBERS=true TAGGING_POLICY=All REMOVE_DUPLICATES=true ASSUME_SORT_ORDER=coordinate READ_NAME_REGEX=null INDEX=true.

### RNA-seq data processing

The quality of the raw RNA-seq data was assessed using FastQC v0.11.9 [101]. The data was aligned in single-end mode against the *Cannabis sativa* reference genome (cv. cs10; accession number GCF_900626175.2) using HISAT2 v2.1.0 (default parameters) [102]. The mapped reads were sorted using Samtools v1.9 and transcript per million (TPM) counts were generated using StringTie v2.1.3b [100, 103]. Gene-level quantification was performed on the mapped reads using featureCounts from the Subread v2.0.0 package [104]. Principal component analysis (PCA) plots were generated in R v4.1 using the normalised read counts from the DESeq2 v1.44.0 R package [105].

### Peak Calling

Narrow peak type data was called for each biological replicate for the narrow peak type H3K4me3 and for the mixed peak type data H3K56ac using model based analysis of ChIP-seq (MACS2 2.2.7) using the parameters bam paired end function -f BAMPE and mappable genome size -g 736579359 -B and a relaxed p value cut off -p 0.1 as indicated for downstream irreproducible discovery rate (IDR) analysis [106]. Statistically significant, replicable peaks were then estimated using the irreproducible discovery rate (IDR 2.0.4.2) [107]. An IDR threshold or false discovery rate of 0.05 was applied for transcription factor-like H3K4me3 narrow peaks.

There was no community standardised approach for handling mixed peak types like H3K56ac. We consequently handled the dataset as a narrow peak type dataset, as we were primarily interested in the narrow domain functionality of H3K56ac [108]. As IDR is considered particularly stringent in calling reproducible peaks for narrow peak type data, and there is inherent broad domain noise in H3K56ac data, we increased the IDR or false discovery rate cutoff threshold to 0.1 for H3K56ac datasets.

The ENCODE consortium does not have stringent guidelines on how to process biological replicates for mixed peak type data, such as H3K27me3 and H2A.Z broad domains. We took a conservative approach to identifying reproducible broad domains between biological replicates for both H2A.Z and H3K27me3. We used a new implementation of the spatial clustering for identification of ChIP-seq regions (SICER), epic2 0.0.52 specifically designed for broad domain peak type data using default parameters. Prior to epic2 calling of broad domains we scaled each replicate and its corresponding input control, as epic2 does not have auto-scaling functionality like MACS2 software, using the deepTools 3.5.1 bamcoverage option and -- scalefactor parameter. We then identified the replicable broad domains between the replicates using BEDTools 2.3.0 intersect option requiring that broad domains share at least 30% reciprocal overlaps -f 0.3 -r.

### Gene and KEGG ontology analysis

To identify genes associated with various chromatin states we used BEDTools 2.3.0 intersect function to find genes genome-wide that overlapped with replicable peaks for each histone mark/modification. We then used these gene lists to perform Gene ontology and Kyoto encyclopedia of genes and genomes (KEGG) ontology analysis. We used the R-Package ClusterProfiler 4.0 [109] to perform gene ontology analysis using an in-house gene ontology database for the cs10 reference assembly (GCF_900626175.2) created using PANNZER2 [110]. Similarly, we used ClusterProfiler 4.0 to perform KEGG pathway enrichment using the available *C. sativa* KEGG ontologies for the cs10 reference assembly (GCF_900626175.2). For glandular trichome specific analysis we used BEDTools intersect -v function to report peaks that only occurred in glandular trichomes compared to stems and with that output we applied the same operation with the leaf dataset to identify glandular trichome specific peaks for each chromatin type. The genes intersecting the glandular trichome specific chromatin peaks were then subject to ontology analysis as above.

### Differential gene expression and gene set enrichment analysis

Differential gene expression was carried out on the RNA-seq datasets using the R-package DESeq2 with default parameters [105]. The DESeq2 output results were then brought forward for gene set enrichment analysis using the ClusterProfiler 4.0 package ridgeplot function using the PANNZER2 cs10 (GCF_900626175.2) gene ontology database.

### Cis-regulatory motif analysis

Glandular trichome specific distal (≥1.5 Kbp from TSS) H3K56ac peaks were determined using the BEDTools 2.3.0 intersect -v function. The nucleotide sequences within peaks were extracted using the BEDTools 2.3.0 getfasta option. Similarly, the BEDTools 2.3.0 getfasta option was used to determine the nucleotide sequences of all glandular trichome H3K56ac peaks which were then used as background control for motif discovery. Motif discovery was carried out using MEME suite tools [111]. Glandular trichome specific H3K56ac chromatin sequences were enriched for motifs using the simple enrichment analysis (SEA) with default settings, cross-referencing the JASPAR CORE (2022) plants non-redundant motif database [111]. The significantly (q < 0.05) motifs were then plotted by transcription factor family vs - log10(Q-value) using the beeswarm R-package.

### Distal enhancer and target prediction leveraging multiomic datasets

Using the list of glandular trichome specific H3K56ac peaks we then determined gene distal peaks occurring ≥1.5 kb from gene TSSs using the hypergeometric optimization of motif enrichment (HOMER) Perl script getDistalPeaks.pl using the parameters -gid -d 1500, then re-run using the parameter-targets to produce a list of the nearest gene-distal H3K56ac peaks [112]. We then used a custom python script to match the list of putative target genes to their corresponding DESeq2 differentially expressed gene ID in glandular trichome vs stem and glandular trichome vs leaf DESeq2 outputs. The list of genes was then filtered to include only those genes that show transcriptomic evidence of strong glandular trichome specific gene expression (log2FC>2; p.adj <0.05) in both trichome vs. stem and trichome vs. leaf datasets.

## Availability of Data and Materials

Chromatin immunoprecipitation sequencing data was made publicly available on the sequencing read archive (SRA) under BioProject ID PRJNA1128358 and complimentary RNA sequencing datasets were registered under BioProject ID PRJNA1128734.

## Acknowledgements

LJC received a PhD scholarship from La Trobe University Graduate Research School. We thank Asha Haslem for providing sequencing services at the La Trobe University Genomics Platform.

## Author Information

### Authors & Affiliations

La Trobe Institute for Sustainable Agriculture and Food, La Trobe University, AgriBio Building, Bundoora, VIC 3086, Australia

Lee James Conneely, Sophia Ng, Muluneh Oli, Bhavna Hurgobin, Mathew Graham Lewsey

Australian Research Council Research Hub for Medicinal Agriculture, La Trobe University, AgriBio Building, Bundoora, VIC 3086, Australia

Lee James Conneely, Sophia Ng, Muluneh Oli, Bhavna Hurgobin, Mathew Graham Lewsey

Australian Research Council Centre of Excellence in Plants for Space, La Trobe University, Bundoora, Victoria, Australia

Lee James Conneely, Mathew Graham Lewsey

## Contributions

L.J.C. and M.G.L. Conceptualised and designed the study. L.J.C. performed the chromatin immunoprecipitation of each histone mark, prepared the sequencing libraries, and performed genomic alignment of the reads. M.O. extracted RNA from each of the samples, S.N. prepared the RNA sequencing libraries, and B.H. performed alignment of the RNA sequencing reads to genome. L.J.C performed normalisation and peak calling of the ChIP-seq reads, differential gene expression of the RNA-seq reads, gene ontology analysis of both RNA-seq and ChIP-seq data, motif discovery enrichment, and integration of RNA-seq and ChIP-seq data. L.J.C. wrote the original manuscript draft and M.G.L. edited the original draft manuscript. L.J.C. wrote the final version. All authors read and approved the final manuscript.

## Corresponding author

Mathew Graham Lewsey

## Ethics Declarations

### Ethics approval and consent to participate

Not applicable.

### Consent for publication

Not applicable.

## Supplementary Information

### Additional file 1

Supporting figures.

### Additional file 2

HOMER peak annotation of each histone modification in the glandular trichome datasets.

### Additional file 3

HOMER peak annotation of each histone modification in the vegetative leaf datasets.

### Additional file 4

HOMER peak annotation of each histone modification in the stem (internode) datasets.

### Additional file 5

Gene ontology analysis of the genes containing each histone modification for glandular trichomes.

### Additional file 6

KEGG pathway enrichment analysis of the genes containing overlapping H3K4me3/H3K27me3 histone modifications in glandular trichomes.

### Additional file 7

Pairwise differential gene expression analysis of glandular trichome, stem (internode), and vegetative leaf RNA-seq datasets.

### Additional file 8

Pairwise gene set enrichment analysis of glandular trichomes, stem (internode), and vegetative leaf RNA-seq datasets.

### Additional file 9

KEGG pathway enrichment analysis of genes associated with histone modification peaks occurring exclusively in glandular trichomes.

### Additional file 10

Motif enrichment of transcription start site distal glandular trichome specific H3K56ac peaks.

### Additional file 11

List of putative glandular trichome specific enhancers.

### Additional file 12

List of buffers used for nuclei isolation and chromatin immunoprecipitation.

