## Additional File 1 for "Characterization of the *Cannabis sativa* glandular trichome epigenome"

### Additional File 1. Supporting Figures and Tables

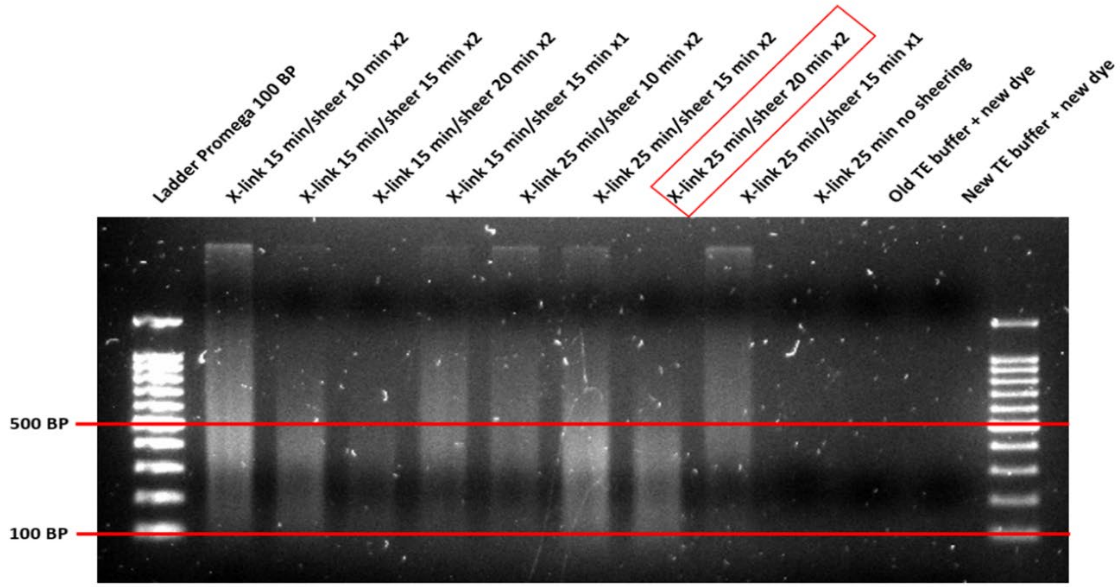


**Figure S1. Optimization of Chromatin fragmentation conditions.** Chromatin from *Cannabis sativa* was subjected to various sonication times and cycles under different cross-linking times to choose the most suitable strategy of obtaining chromatin fragments in the target 100 – 500 bp range. A final cross-linking time of 25 mins in combination with 2 cycles of 20 min sheering on high setting was selected for downstream immunoprecipitation.

**Table S1. Qubit dsDNA high sensitivity Assay results of immunoprecipitated chromatin from glandular trichomes, stalks, and vegetative leaves.** dsDNA content was estimated for each Immunoprecipitate to ensure a minimum sufficient mass of >10 pg was available for downstream library preparation using the Swift ACCEL-NGS® 2S Plus DNA library prep kit.

| Sample ID | dsDNA concentration | |
|---|---|---|
| | Biological Replicate 1 | Biological Replicate 2 |
| INPUT control (Trichomes) | 8.96 ng/µL | 4.22 ng/µL |
| INPUT control (Stalks) | 3.22 ng/µL | 3.06 ng/µL |
| INPUT control (Vegetative leaves) | 4.7 ng/µL | 4.90 ng/µL |
| H2A.Z (Trichomes) | 0.014 ng/µL | 0.0112 ng/µL |
| H2A.Z (Stalks) | 0.0062 ng/µL | 0.0066 ng/µL |
| H2A.Z (Vegetative leaves) | 0.0051 ng/µL | 0.0104 ng/µL |
| H3K4me3 (Trichomes) | 0.14 ng/µL | 0.0504 ng/µL |
| H3K4me3 (Stalks) | 0.0850 ng/µL | 0.0804 ng/µL |
| H3K4me3 (Vegetative leaves) | 0.0574 ng/µL | 0.0240 ng/µL |
| H3K56ac (Trichomes) | 0.112 ng/µL | 0.0778 ng/µL |
| H3K56ac (Stalks) | 0.0188 ng/µL | 0.0162 ng/µL |
| H3K56ac (Vegetative leaves) | 0.0148 ng/µL | 0.0148 ng/µL |
| H3K27me3 (Trichomes) | 0.0062 ng/µL | 0.0069 ng/µL |
| H3K27me3 (Stalks) | 0.0108 ng/µL | 0.0106 ng/µL |
| H3K27me3 (Vegetative leaves) | 0.0050 ng/µL | 0.0071 ng/µL |

**INPUT Control Trichomes Rep. 1**

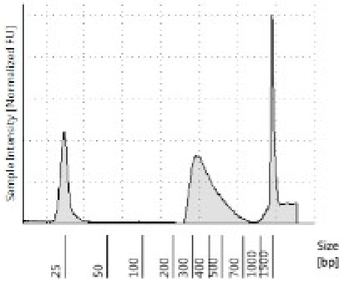


**INPUT Control Stem (internode) Rep. 1**

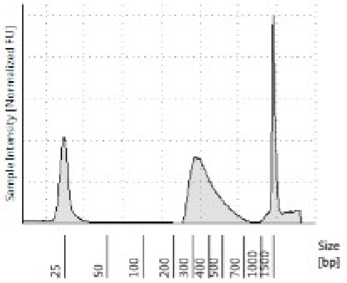


**INPUT control Vegetative Leaves Rep. 1**

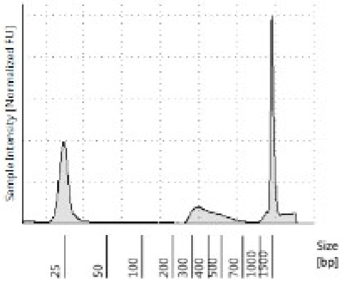


**H2A.Z trichomes Rep. 1**

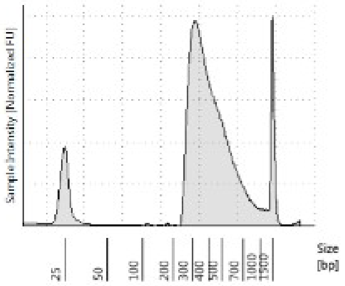


**H2A.Z Stem (internode) Rep. 1**

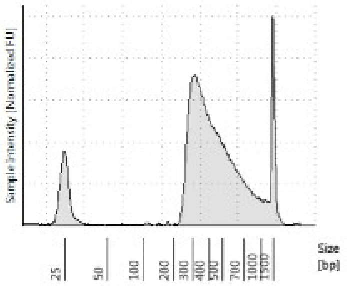


**H2A.Z Vegetative leaves Rep. 1**

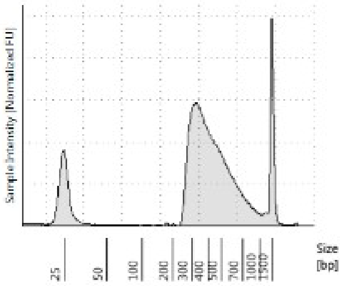


**H3K4me3 Trichomes Rep. 1**

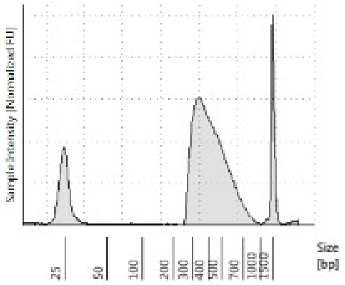


**H3K4me3 Stem (internode) Rep. 1**

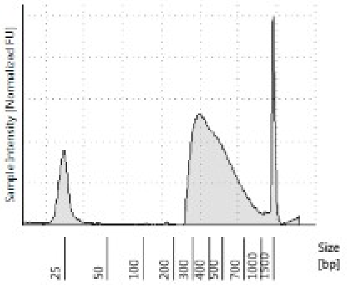


**H3K4me3 Vegetative leaves Rep. 1**

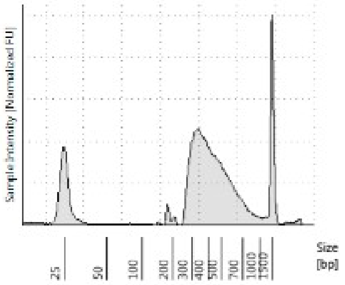


**H3K56ac Trichomes Rep. 1**

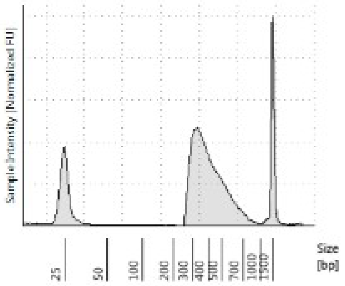


**H3K56ac Stem (internode) Rep. 1**

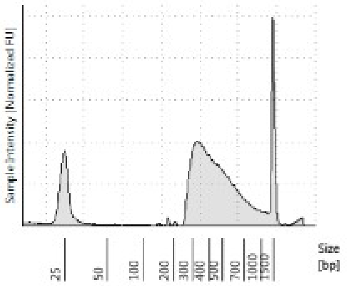


**H3K56ac Vegetative leaves Rep. 1**

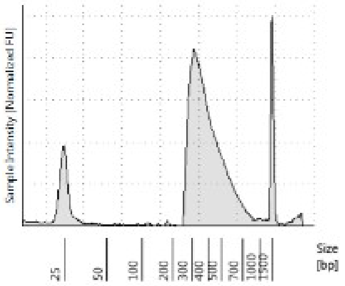


**H3K27me3 Trichomes Rep. 1**

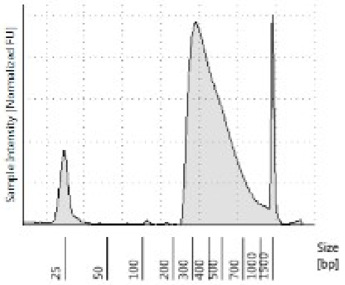


**H3K27me3 Stem (internode) Rep. 1**

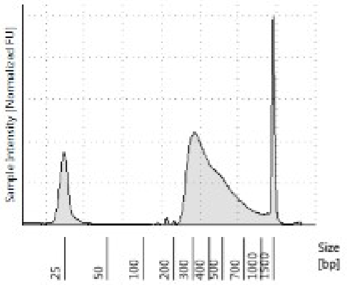


**H3K27me3 Vegetative leaves Rep. 1**

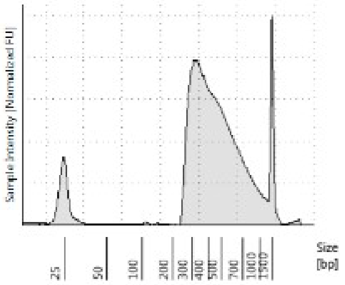

**INPUT Control Trichomes Rep. 2**

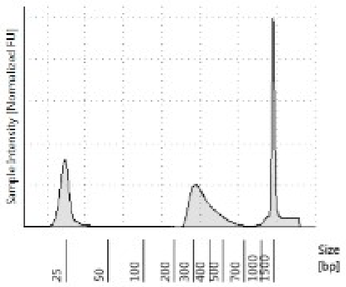


**INPUT Control Stem (internode) Rep. 2**

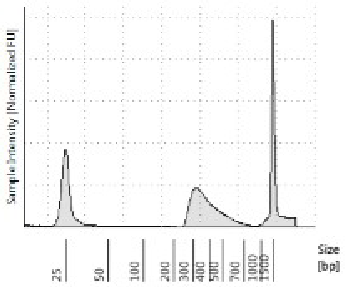


**INPUT control Vegetative Leaves Rep. 2**

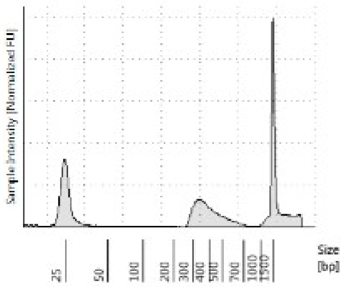


**H2A.Z trichomes Rep. 2**

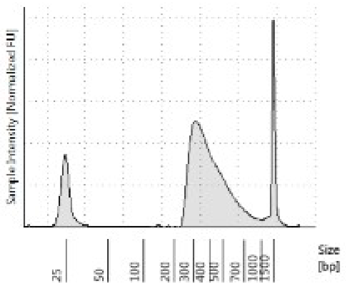


**H2A.Z Stem (internode) Rep. 2**

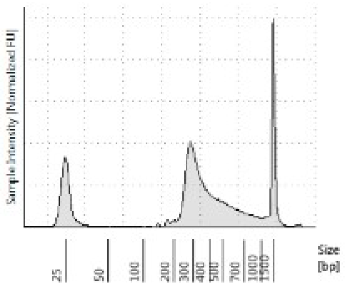


**H2A.Z Vegetative leaves Rep. 2**

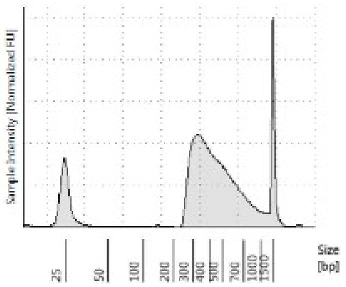


**H3K4me3 Trichomes Rep. 2**

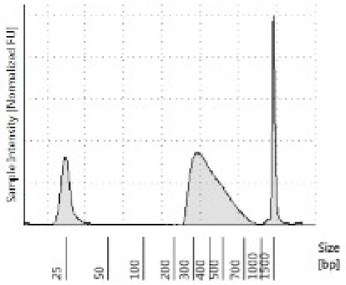


**H3K4me3 Stem (internode) Rep. 2**

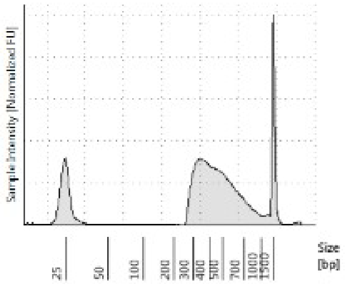


**H3K4me3 Vegetative leaves Rep. 2**

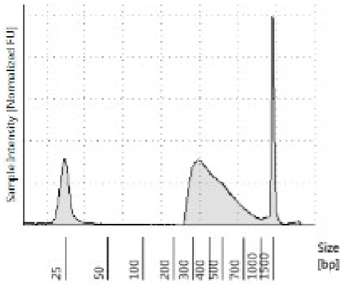


**H3K56ac Trichomes Rep. 2**

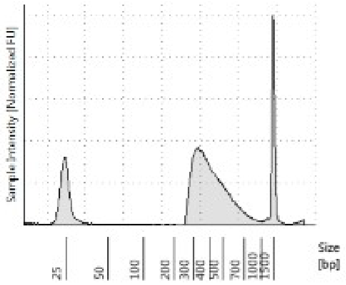


**H3K56ac Stem (internode) Rep. 2**

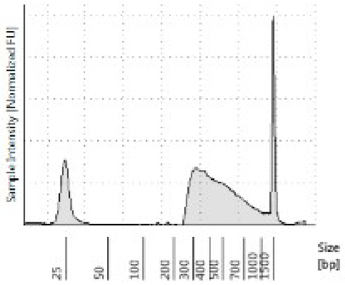


**H3K56ac Vegetative leaves Rep. 2**

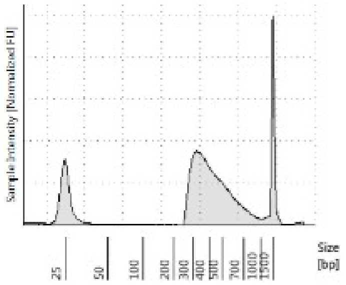


**H3K27me3 Trichomes Rep. 2**

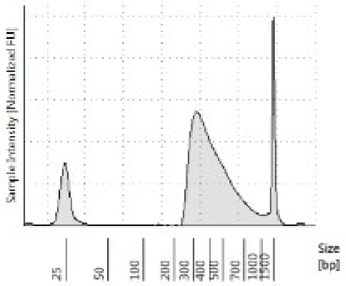


**H3K27me3 Stem (internode) Rep. 2**

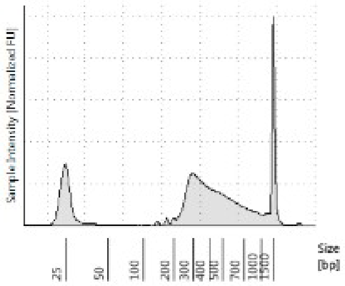


**H3K27me3 Vegetative leaves Rep. 2**

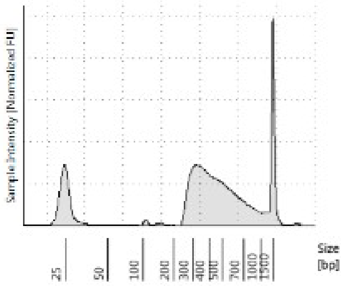

**Figure S2. 2200 Tapestation D1000 electropherograms of ChIP-seq libraries.** 2200 Tapestation D1000 electropherograms of 30 ChIP-seq libraries including INPUT control and 4 histone marks H2A.Z, H3K4me3, H3K56ac, and H3K27me3 for *Cannabis sativa* glandular trichomes, stem (internode), and vegetative leaves in biological duplicate.

**Table S2. ChIP-seq libraries fragment size and concentration estimation.** Average fragment size estimation from 2200 Tapestation D1000 results, and qubit concentration estimates for each library. Codes for Swift biosciences Accel-NGS 2S indexing adapters used for each library listed under the indexing adapters as well as the corresponding plate position.

| Sample ID | Indexing adapter | Plate position | Average Fragment length bp | dsDNA ng/uL |
|---|---|---|---|---|
| INPUT control Trichomes Rep. 1 | U037 | A1 | 402 | 61.6 |
| INPUT control Stem (internode) Rep. 1 | U038 | A2 | 393 | 59 |
| INPUT control Vegetative leaf Rep. 1 | U039 | A3 | 422 | 62.2 |
| H2A.Z Trichomes Rep. 1 | U040 | A4 | 430 | 67.4 |
| H2A.Z Stem (internode) Rep. 1 | U041 | A5 | 452 | 52.2 |
| H2A.Z Vegetative leaf Rep. 1 | U042 | A6 | 444 | 71.4 |
| H3K4me3 Trichomes Rep. 1 | U043 | A7 | 431 | 80.2 |
| H3K4me3 Stem (internode) Rep. 1 | U044 | A8 | 462 | 72.4 |
| H3K4me3 Vegetative leaf Rep. 1 | U045 | A9 | 439 | 66.6 |
| H3K56ac Trichomes Rep. 1 | U046 | A10 | 418 | 79.6 |
| H3K56ac Stem (internode) Rep. 1 | U047 | A11 | 464 | 61 |
| H3K56ac Vegetative leaf Rep. 1 | U048 | A12 | 410 | 71 |
| H3K27me3 Trichomes Rep. 1 | U049 | B1 | 443 | 67.2 |
| H3K27me3 Stem (internode) Rep. 1 | U050 | B2 | 434 | 58.2 |
| H3K27me3 Vegetative leaf Rep. 1 | U051 | B3 | 470 | 71.5 |
| INPUT control Trichomes Rep. 2 | U061 | G1 | 377 | 55.4 |
| INPUT control Stem (internode) Rep. 2 | U062 | G2 | 319 | 57.2 |
| INPUT control Vegetative leaf Rep. 2 | U063 | G3 | 410 | 48.8 |
| H2A.Z Trichomes Rep. 2 | U064 | G4 | 420 | 53.4 |
| H2A.Z Stem (internode) Rep. 2 | U065 | G5 | 410 | 45.2 |
| H2A.Z Vegetative leaf Rep. 2 | U066 | G6 | 463 | 68.2 |
| H3K4me3 Trichomes Rep. 2 | U067 | G7 | 415 | 60.2 |
| H3K4me3 Stem (internode) Rep. 2 | U068 | G8 | 476 | 70.4 |
| H3K4me3 Vegetative leaf Rep. 2 | U069 | G9 | 450 | 66.0 |
| H3K56ac Trichomes Rep. 2 | U070 | G10 | 429 | 62.2 |
| H3K56ac Stem (internode) Rep. 2 | U071 | G11 | 478 | 60.4 |
| H3K56ac Vegetative leaf Rep. 2 | U072 | G12 | 442 | 71.4 |
| H3K27me3 Trichomes Rep. 2 | U073 | H1 | 436 | 65.2 |
| H3K27me3 Stem (internode) Rep. 2 | U074 | H2 | 448 | 40.4 |
| H3K27me3 Vegetative leaf Rep. 2 | U075 | H3 | 466 | 51.2 |

**Table S3a. Bowtie2 Alignment mapping statistics of paired end ChIP-seq data alignment to cs10 (GCF_900626175.2).** Table of the mapping statistics for each ChIP-seq library in this study.

| Sample | Total reads | Overall alignment rate | Unmapped Reads | Uniquely mapped reads | Multi-mapping reads |
|---|---|---|---|---|---|
| INPUT control Trichomes Rep. 1 | 25,157,536 | 90.78% | 15.11% | 28.31% | 56.59% |
| INPUT control Stem (internode) Rep. 1 | 21,493,669 | 91.74% | 13.80% | 26.23% | 59.97% |
| INPUT control Vegetative leaf Rep. 1 | 26,893,660 | 91.47% | 14.93% | 26.03% | 59.04% |
| H2A.Z Trichomes Rep. 1 | 12,680,890 | 82.71% | 22.69% | 23.60% | 53.71% |
| H2A.Z Stem (internode) Rep. 1 | 13,028,860 | 65.50% | 39.22% | 18.27% | 42.50% |
| H2A.Z Vegetative leaf Rep. 1 | 18,266,832 | 86.91% | 18.83% | 24.37% | 56.80% |
| H3K4me3 Trichomes Rep. 1 | 28,609,394 | 89.27% | 15.10% | 61.42% | 23.49% |
| H3K4me3 Stem (internode) Rep. 1 | 15,347,697 | 89.83% | 15.57% | 51.88% | 32.55% |
| H3K4me3 Vegetative leaf Rep. 1 | 11,303,989 | 87.75% | 17.95% | 36.51% | 45.54% |
| H3K56ac Trichomes Rep. 1 | 10,440,217 | 89.65% | 16.37% | 29.64% | 53.99% |
| H3K56ac Stem (internode) Rep. 1 | 10,513,921 | 85.66% | 20.23% | 29.14% | 50.63% |
| H3K56ac Vegetative leaf Rep. 1 | 11,569,419 | 90.44% | 15.14% | 25.07% | 59.79% |
| H3K27me3 Trichomes Rep. 1 | 15,697,513 | 83.17% | 22.87% | 26.37% | 50.76% |
| H3K27me3 Stem (internode) Rep. 1 | 14,592,876 | 83.73% | 22.58% | 24.95% | 52.47% |
| H3K27me3 Vegetative leaf Rep. 1 | 10,488,852 | 79.78% | 26.06% | 23.18% | 50.76% |
| INPUT control Trichomes Rep. 2 | 26,716,190 | 90.85% | 14.89% | 28.07% | 57.04% |
| INPUT control Stem (internode) Rep. 2 | 20,994,779 | 91.71% | 14.00% | 25.80% | 60.19% |
| INPUT control Vegetative leaf Rep. 2 | 31,365,486 | 91.45% | 14.51% | 26.30% | 59.19% |
| H2A.Z Trichomes Rep. 2 | 14,816,661 | 81.30% | 23.83% | 22.86% | 53.31% |
| H2A.Z Stem (internode) Rep. 2 | 13,879,251 | 41.55% | 61.47% | 11.69% | 26.84% |
| H2A.Z Vegetative leaf Rep. 2 | 13,114,965 | 80.28% | 25.29% | 22.13% | 52.58% |
| H3K4me3 Trichomes Rep. 2 | 20,846,135 | 77.82% | 25.62% | 55.28% | 19.11% |
| H3K4me3 Stem (internode) Rep. 2 | 11,454,387 | 89.27% | 16.10% | 50.85% | 33.05% |
| H3K4me3 Vegetative leaf Rep. 2 | 13,593,404 | 90.10% | 15.21% | 46.31% | 38.48% |
| H3K56ac Trichomes Rep. 2 | 17,838,025 | 89.46% | 16.49% | 30.16% | 53.35% |
| H3K56ac Stem (internode) Rep. 2 | 12,534,003 | 79.36% | 26.17% | 30.05% | 43.78% |
| H3K56ac Vegetative leaf Rep. 2 | 17,122,515 | 90.14% | 15.68% | 26.27% | 58.05% |
| H3K27me3 Trichomes Rep. 2 | 14,239,146 | 83.91% | 21.83% | 25.77% | 52.40% |
| H3K27me3 Stem (internode) Rep. 2 | 10,228,339 | 66.16% | 38.75% | 20.78% | 40.48% |
| H3K27me3 Vegetative leaf Rep. 2 | 13,828,751 | 86.62% | 19.64% | 25.77% | 54.60% |

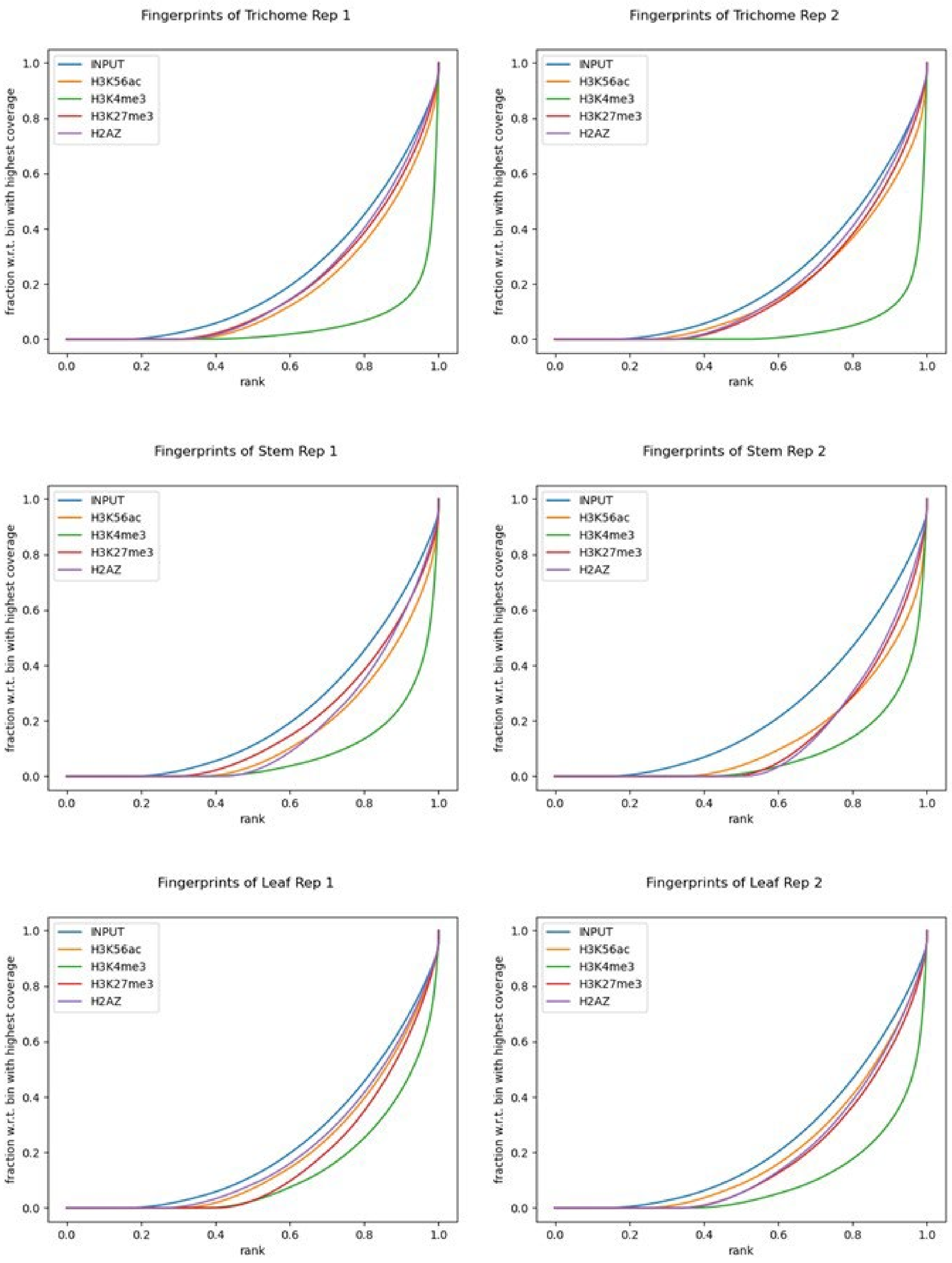


**Figure S3. Fingerprint plots of INPUT control and histone marks for each tissue type.** Fingerprint plots showing how the sequencing reads of each histone mark distribute across the *C. sativa* genome. Narrow peak data, H3K4me3 in this instance, tend toward acute right handed angles whereby the majority of reads are distributed in a small proportion of the genome and broad histone marks such as H3K56ac, H3K27me3 as well as the histone variant H2A.Z indicate a comparatively linear distribution throughout the genome similar to, yet discriminate from, the distribution of reads observed in the INPUT control

**Table S3b. HISAT2 Alignment mapping statistics of single end RNA sequencing data alignment to cs10 (GCF_900626175.2).**

| Sample | No. of raw reads | No. of mapped reads | %mapped reads |
|---|---|---|---|
| Leaf Rep. 1 | 23,473,287 | 21,208,394 | 90.35 |
| Leaf Rep. 2 | 24,274,197 | 21,936,259 | 90.37 |
| Leaf Rep. 3 | 23,267,045 | 20,984,219 | 90.19 |
| Shoot Rep. 1 | 27,917,083 | 25,264,096 | 90.50 |
| Shoot Rep. 2 | 19,416,942 | 17,485,856 | 90.05 |
| Shoot Rep. 3 | 25,321,091 | 22,916,816 | 90.50 |
| Trichomes Rep. 1 | 29,914,474 | 26,944,383 | 90.07 |
| Trichomes Rep. 2 | 26,542,737 | 24,012,723 | 90.47 |
| Trichomes Rep. 3 | 25,246,050 | 22,817,394 | 90.38 |

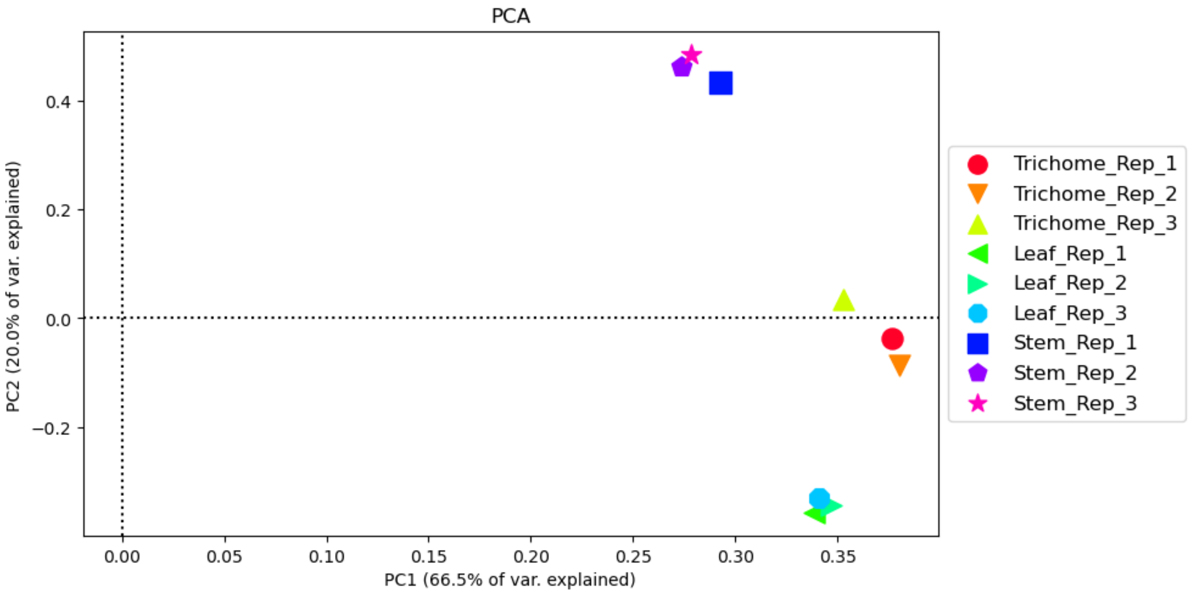


**Figure S4. Principal component analysis of RNA sequencing data from glandular trichomes, Stem (internode), and leaf tissues in triplicate.** PCA indicates agreement between the biological replicates and distinction between the sample types. 66.5 % of the variance can be resolved by component 1.

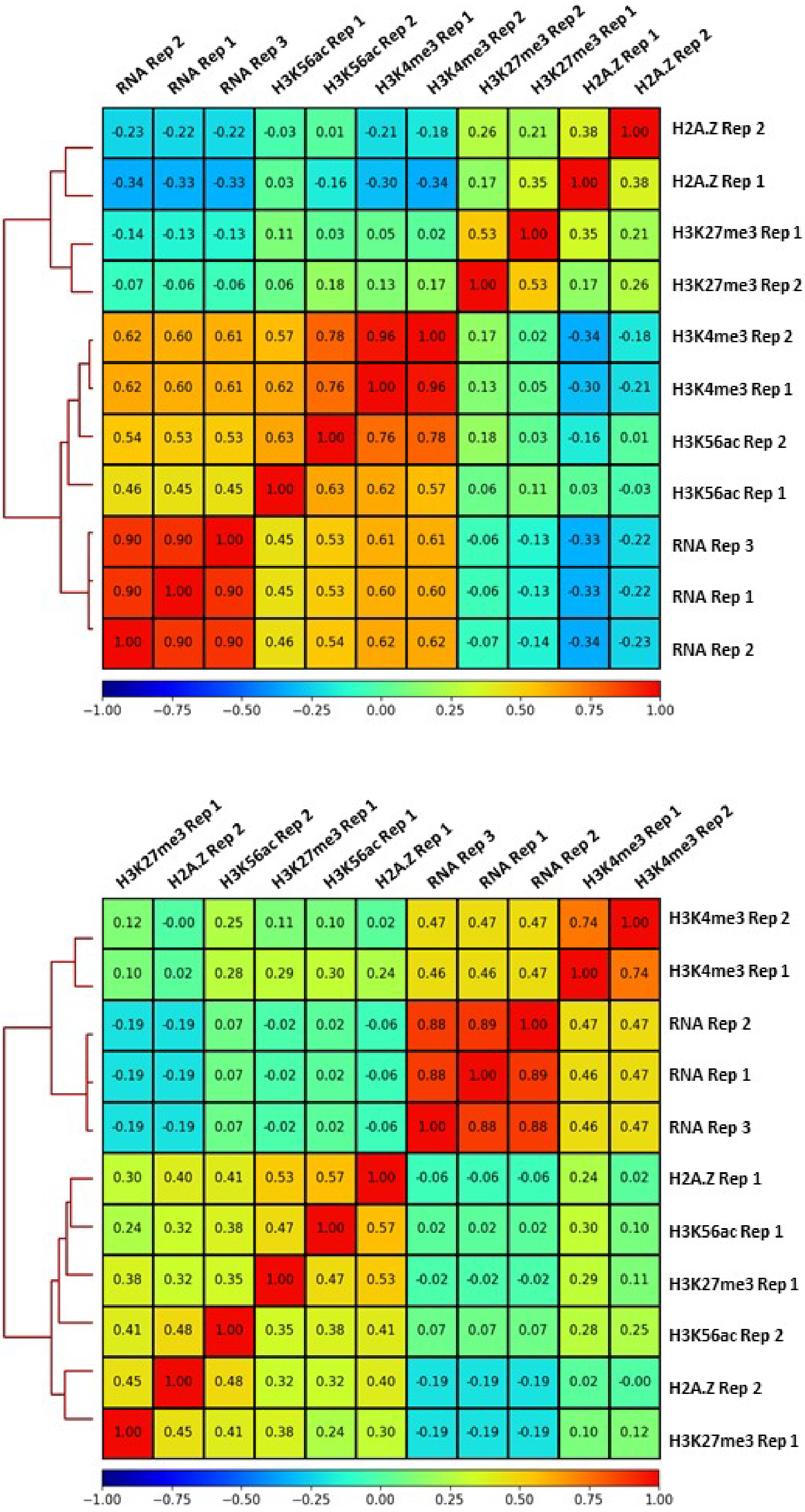


**Figure S5. Spearman correlation plots.** Data relationships between RNA seq and Histone marks were analysed by spearman correlation and plotted for stem (top) and leaf (bottom). These figures are the stem and leaf compliment of Fig. 1b.

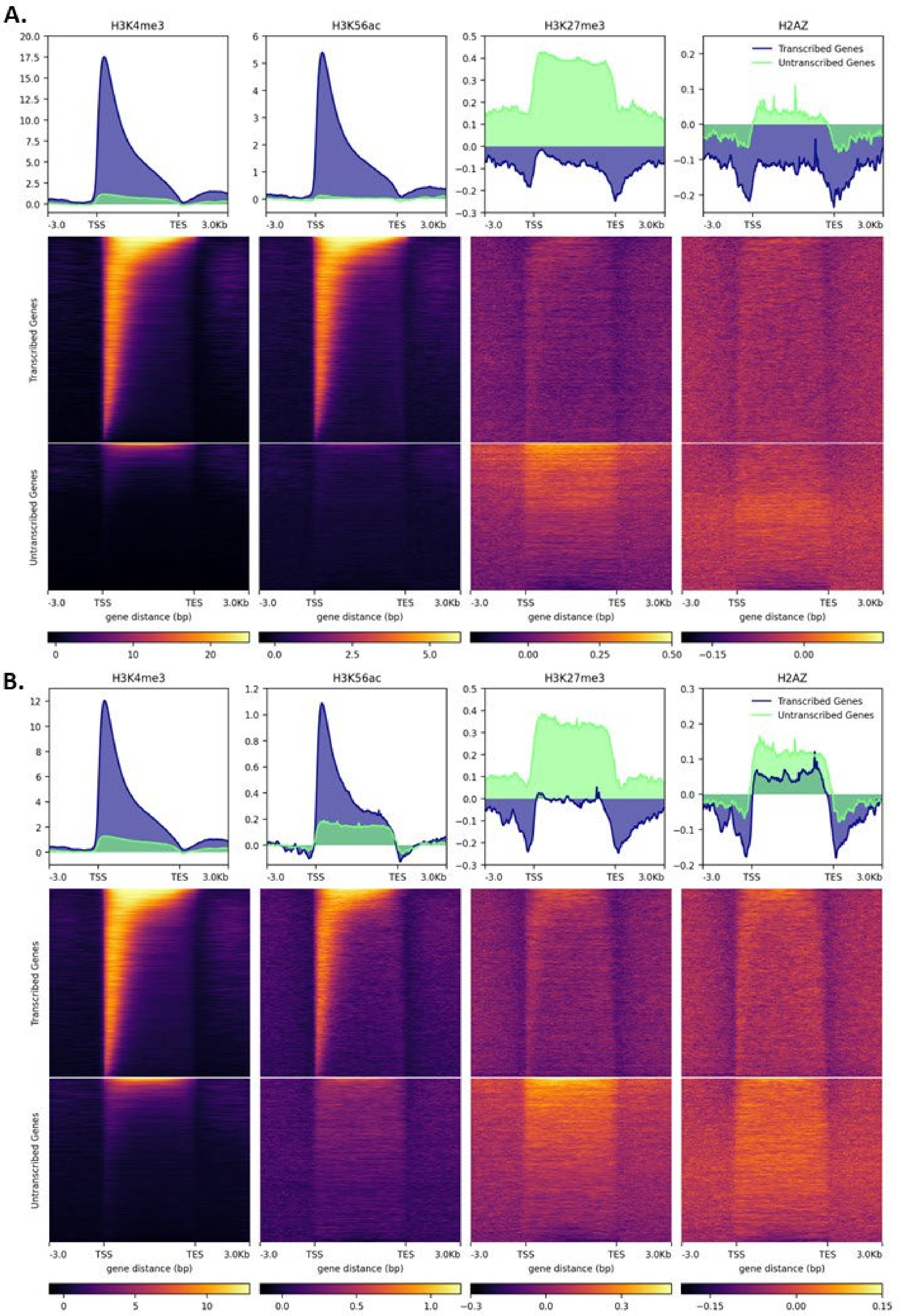


**Figure S6. Integrated expression TSS plots of histone marks and variant.** Histone mark relationship and distribution in genes transcribed and untranscribed in that respective tissue. Stem (A) and leaf (B). Compliment of trichomes Fig. 1e.

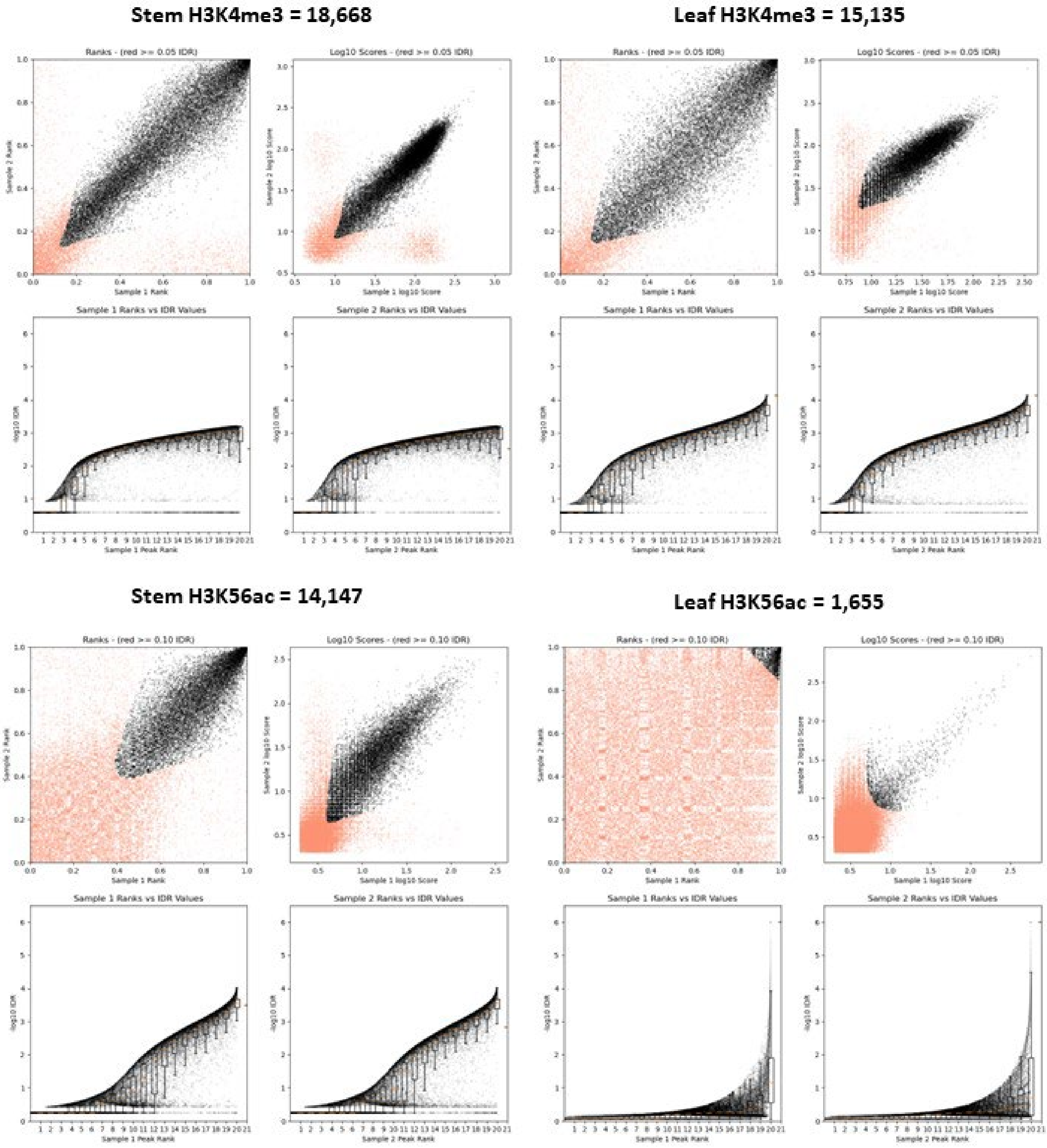


**Figure S7. Irreproducible discovery rate analysis of Stem and Leaf.** Application of the irreproducible discovery rate analysis method to determine consensus peaks between replicates for both narrow peak H3K4me3 and Mixed peak type H3K56ac replicates for both stem and leaf tissue. Consensus peaks identified are displayed above each panel of four. These are results are the stem and leaf compliment to trichomes found in Fig. 2 a & b.

```
# Number of query intervals (a) H3K4me3 Trichomes: 18774

# Number of db intervals (b) H3K56ac Trichomes: 13449

# Number of overlaps: 9668

# Number of possible intervals (estimated): 333375

# phyper(9668 - 1, 18774, 333375 - 18774, 13449, lower.tail=F)

# Contingency Table Of Counts

#_________________________________________

#           |  in -b       | not in -b    |

#     in -a | 9668         | 9106         |

# not in -a | 3781         | 310820       |

#_________________________________________

# p-values for fisher's exact test

left	right	two-tail	ratio

1	0	0	87.279
```

**Figure S7b. Fisher exact test results H3K4me3 H3K56ac Trichomes.** Null hypothesis is rejected, H3K4me3 and H3K56ac peaks co-occur more than is expected by chance (two-tail p-value < $1x10^{-5}$). With at least 50% reciprocal overlap of peaks.

```
# Number of query intervals: 12601

# Number of db intervals: 10028

# Number of overlaps: 3169

# Number of possible intervals (estimated): 108988

# phyper(3169 - 1, 12601, 108988 - 12601, 10028, lower.tail=F)

# Contingency Table Of Counts

#_________________________________________

#           |  in -b       | not in -b    |

#     in -a | 3169         | 9432         |

# not in -a | 6859         | 89528        |

#_________________________________________

# p-values for fisher's exact test

left	right	two-tail	ratio

1	0	0	4.385
```

**Figure S7c. Fisher exact test results H3K27me3 H2AZ Trichomes.** Null hypothesis is rejected, H3K27me3 and H2AZ peaks co-occur more than is expected by chance (two-tail p-value $< 1x10^{-5}$). With at least 50% reciprocal overlap of peaks.

```
# Number of query intervals: 18774

# Number of db intervals: 10028

# Number of overlaps: 238

# Number of possible intervals (estimated): 137167

# phyper(238 - 1, 18774, 137167 - 18774, 10028, lower.tail=F)

# Contingency Table Of Counts

#_________________________________________

#           |  in -b       | not in -b    |

#     in -a | 238          | 18536        |

# not in -a | 9790         | 108603       |

#_________________________________________

# p-values for fisher's exact test

left    right   two-tail   ratio

0       1       0          0.142
```

**Figure S7d. Fisher exact test results H3K4me3 H3K27me3 Trichomes.** Null hypothesis is rejected, H3K4me3 and H3K27me3 peaks co-occur more than is expected by chance (two-tail p-value $< 1x10^{-5}$). With at least 50% reciprocal overlap of peaks.

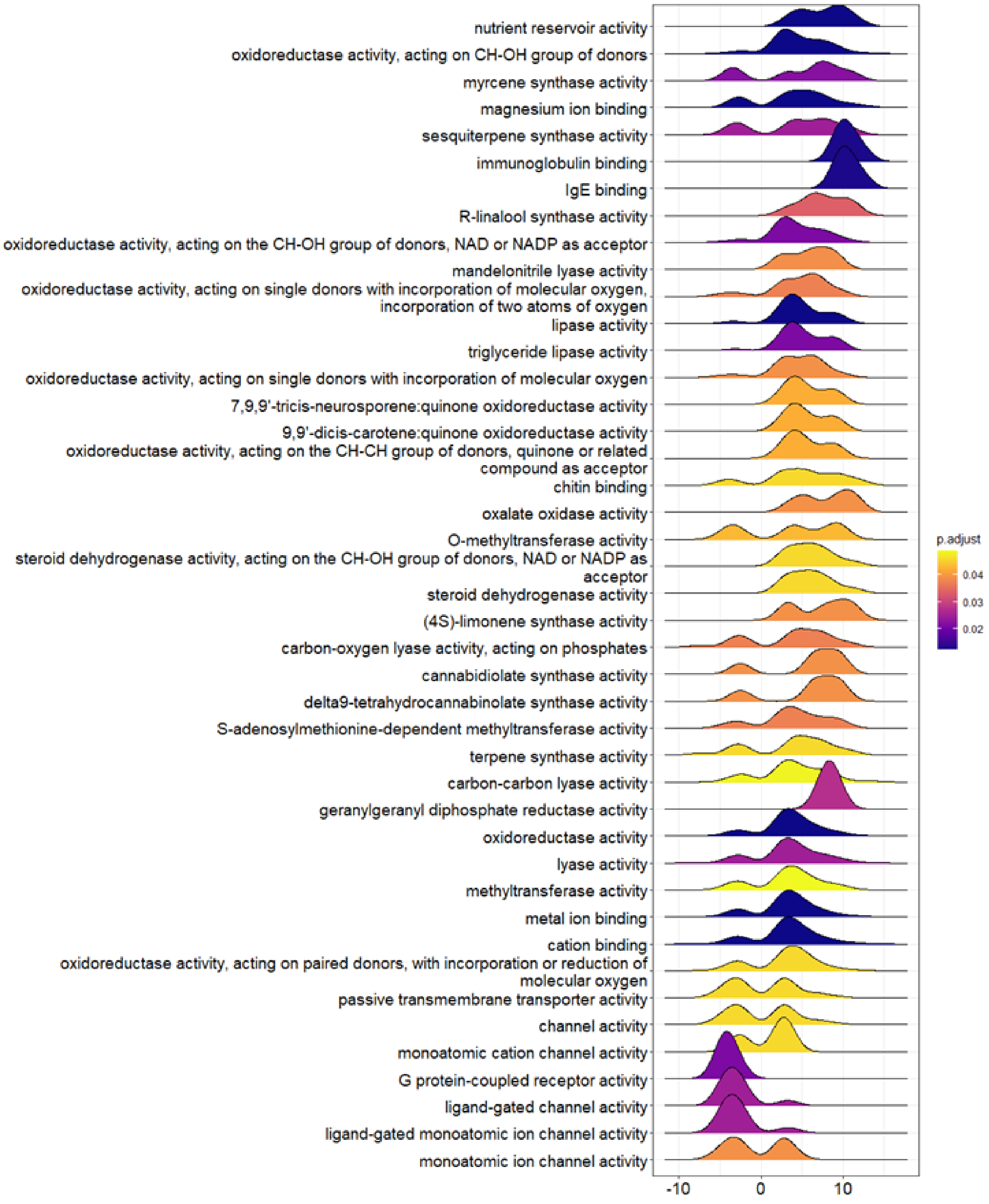


**Figure S8a. Ridge plot of glandular trichome enriched gene sets versus leaf.** Gene set enrichment analysis of glandular trichomes versus leaf molecular function gene ontologies conveyed by means of a ridge plot. Molecular function ontologies are listed and glandular trichome enriched ontologies are denoted by positive integers on the x-axis.

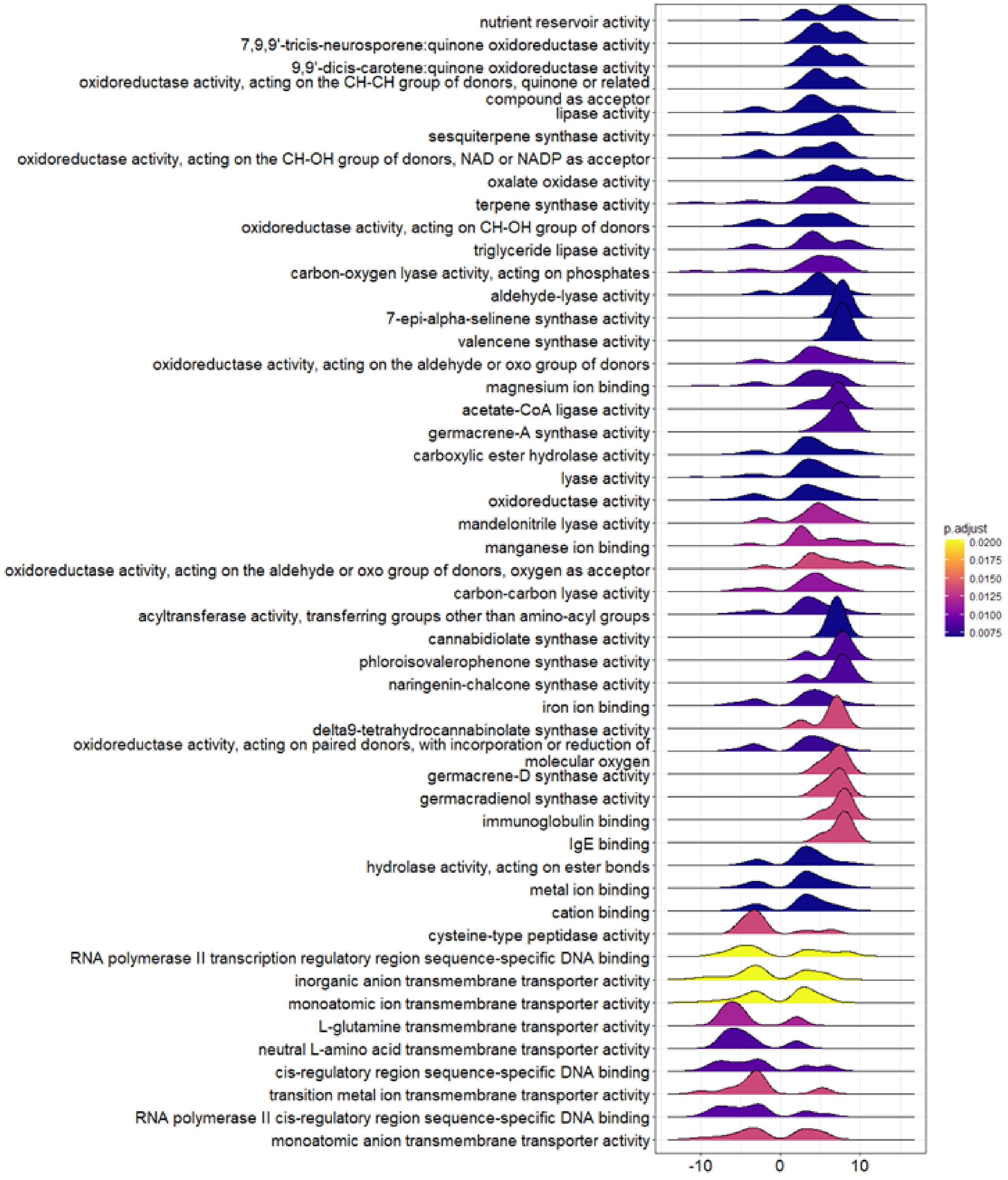


**Figure S8b. Ridge plot of glandular trichome enriched gene sets versus stem.** Gene set enrichment analysis of glandular trichomes versus stem molecular function gene ontologies conveyed by means of a ridge plot. Molecular function ontologies are listed and glandular trichome enriched ontologies are denoted by positive integers on the x-axis.

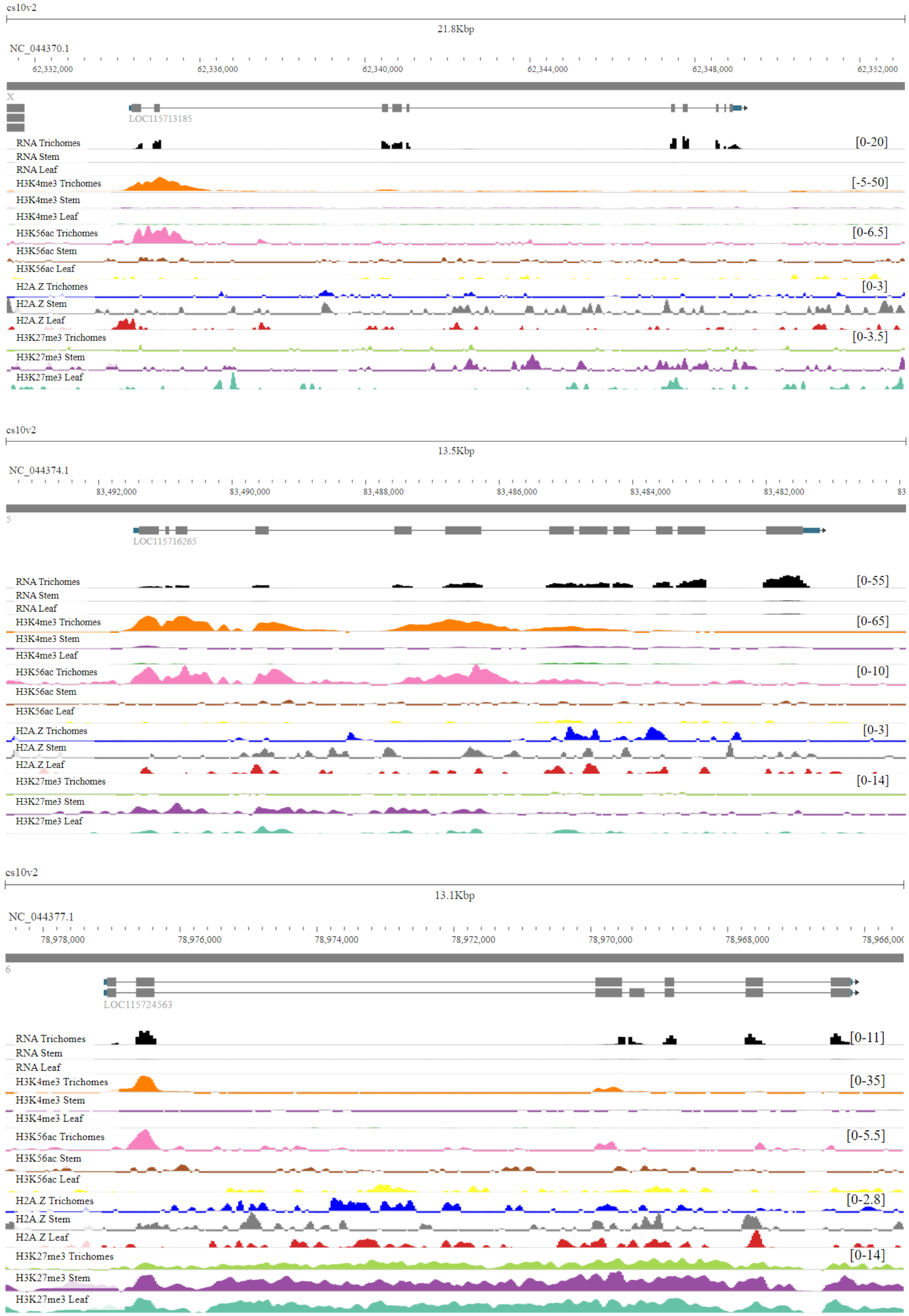
cs10v2
21.8Kbp
NC_044370.1
62,332,000
62,336,000
62,340,000
62,344,000
62,348,000
62,352,000
LOC115713185
RNA Trichomes
RNA Stem
RNA Leaf
H3K4me3 Trichomes
H3K4me3 Stem
H3K4me3 Leaf
H3K56ac Trichomes
H3K56ac Stem
H3K56ac Leaf
H2A.Z Trichomes
H2A.Z Stem
H2A.Z Leaf
H3K27me3 Trichomes
H3K27me3 Stem
H3K27me3 Leaf
[0-20]
[-5-50]
[0-6.5]
[0-3]
[0-3.5]
cs10v2
13.5Kbp
NC_044374.1
83,492,000
83,490,000
83,488,000
83,486,000
83,484,000
83,482,000
LOC115716265
RNA Trichomes
RNA Stem
RNA Leaf
H3K4me3 Trichomes
H3K4me3 Stem
H3K4me3 Leaf
H3K56ac Trichomes
H3K56ac Stem
H3K56ac Leaf
H2A.Z Trichomes
H2A.Z Stem
H2A.Z Leaf
H3K27me3 Trichomes
H3K27me3 Stem
H3K27me3 Leaf
[0-55]
[0-65]
[0-10]
[0-3]
[0-14]
cs10v2
13.1Kbp
NC_044377.1
78,978,000
78,976,000
78,974,000
78,972,000
78,970,000
78,968,000
LOC115724563
RNA Trichomes
RNA Stem
RNA Leaf
H3K4me3 Trichomes
H3K4me3 Stem
H3K4me3 Leaf
H3K56ac Trichomes
H3K56ac Stem
H3K56ac Leaf
H2A.Z Trichomes
H2A.Z Stem
H2A.Z Leaf
H3K27me3 Trichomes
H3K27me3 Stem
H3K27me3 Leaf
[0-11]
[0-35]
[0-5.5]
[0-2.8]
[0-14]

**Figure S9a. Three known trichome genes showing tissue-specific expression and chromatin states.** LOC115713185 (CBGAS), LOC115716265 (ABC transporter B family member 2), and LOC115724563 (alpha-humulene synthase).

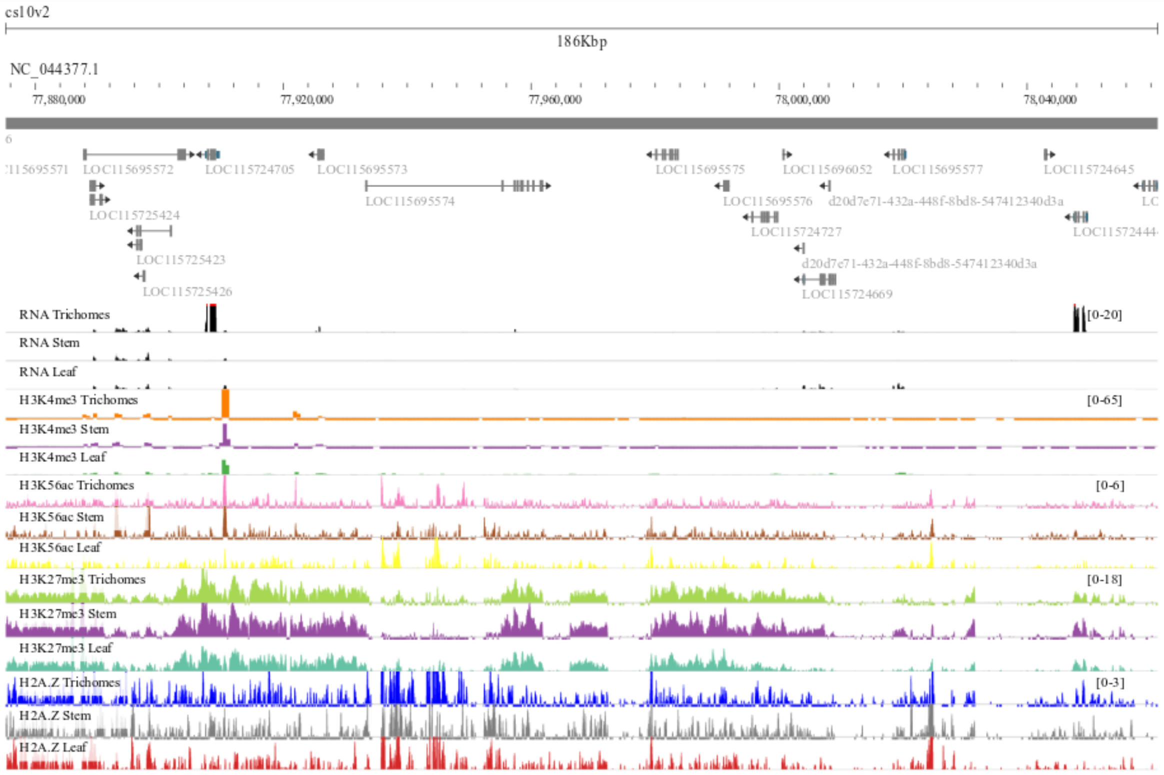


**Figure S9b. Biosynthetic gene cluster showing tissue-specific expression.** Cluster No. 28 biosynthetic gene cluster that contains genes that play role in plant biotic stress resistance to insects and fungus.
