## Additional File 12 for "Characterization of the *Cannabis sativa* glandular trichome epigenome"

Additional File 12. ChIP-seq buffer recipes

Table A1. Nuclear isolation/ cross-linking buffer

| **Nuclear isolation buffer** | **For 120 mL** |
| --- | --- |
| 60mM HEPES pH 8.0 | 7.2 mL 1M HEPES pH 8.0 |
| 1M Sucrose | 60 mL 2M Sucrose (make fresh) |
| 5mM KCl | 0.6 mL 1M KCl |
| 5mM MgCl_2_ | 0.6 mL 1M MgCl_2_ |
| 5mM EDTA pH 8 | 1.2 mL 0.5M EDTA pH 8 |
| 0.6% (v/v) Triton X-100 | 7.2 mL 10% (v/v) Triton X-100 |
| ddH_2_O | 43.2 mL ddH_2_O |
| 0.1-1mM PMSF | 1.2 mL 100mM PMSF (0.1M in ethanol Sigma-Aldrich 93482-50ML-F) |
| 1mM Pepstatin A | 120 µL 1M Pepstatin A (≥90% HPLC Sigma-Aldrich P5318-5MG) |
| Mini-complete (1 mini tablet/10mL) | 12 mini complete tabs (Sigma-Aldrich 11836170001) |

Table A2. Extraction buffer

| **Extraction buffer 2 (4°C)** | **For 10 mL** |
| --- | --- |
| 0.25M Sucrose | 1.25 mL 2M Sucrose |
| 10mM Tris-HCl pH 8.0 | 100 µL 1M Tris-HCl pH 8.0 |
| 10mM MgCl_2_ | 100 µL 1M MgCl_2_ |
| 1% (v/v) Triton X-100 | 1 mL 10% (v/v) Triton X-100 |
| 1mM EDTA pH 8.0 | 20 µL 0.5M EDTA pH 8.0 |
| ddH_2_O | 7.53 mL ddH_2_O |
| 5mM BME | 3.5 µL 14.3M β-mercaptoethanol (BME) |
| 0.1-1mM PMSF | 100 µL 100mM PMSF (Sigma-Aldrich 93482-50ML-F in ethanol) |
| 1mM Pepstatin A | 10 µL 1M Pepstatin A (≥90% HPLC Sigma-Aldrich P5318-5MG) |
| Mini-complete (1 mini tablet/10mL) | 1 mini complete EDTA-free Protease Inhibitor Cocktail tablet (Sigma-Aldrich 11836170001) |

Table A3. Nuclei lysis buffer

| **Nuclei lysis buffer (4°C)** | **For 5 mL** |
| --- | --- |
| 50mM Tris-HCl, pH 8.0 | 250 µL 1M Tris-HCl, pH 8.0 |
| 10mM EDTA pH 8.0 | 100 µL 0.5M EDTA pH 8.0 |
| 1% (w/v) SDS | 500 µL 10% (w/v) SDS |
| ddH_2_O | ddH_2_O to volume |
| 0.1-1mM PMSF | 50 µL 100mM PMSF (Sigma-Aldrich 93482-50ML-F) |
| 1mM Pepstatin A | 5 µL 1M Pepstatin A (≥90% HPLC Sigma-Aldrich P5318-5MG) |
| Mini-complete (1 mini tablet/10mL) | 1/2 mini complete EDTA-free Protease Inhibitor Cocktail tablet (Sigma-Aldrich 11836170001) |

Table A4. ChIP dilution buffer

| **ChIP dilution buffer (4°C)** | **For 40 mL** |
| --- | --- |
| 1.1% (v/v) Triton X-100 | 4.4 mL 10% (v/v) Triton X-100 |
| 1.2mM EDTA pH 8.0 | 96 µL 0.5M EDTA pH 8.0 |
| 16.7mM Tris-HCl, pH 8.0 | 668 µL 1M Tris-HCl, pH 8.0 |
| 167mM NaCl | 1.336 mL 5M NaCl |
| ddH_2_O | ddH_2_O to volume |
| 0.1-1mM PMSF | 400 µL 100mM PMSF (Sigma-Aldrich 93482-50ML-F) |
| 1mM Pepstatin A | 40 µL 1M Pepstatin A (≥90% HPLC Sigma-Aldrich P5318-5MG) |
| Mini-complete (1 mini tablet/10mL) | 4 mini complete EDTA-free Protease Inhibitor Cocktail tablets (Sigma-Aldrich 11836170001) |

Table A5. Low salt wash buffer

| **Low salt wash buffer (4°C)** | **50 mL Low salt wash buffer** |
| --- | --- |
| 150mM NaCl | 1.5 mL 5M NaCl |
| 0.1% (w/v) SDS | 0.5 mL 10% (w/v) SDS |
| 1% (v/v) TritonX-100 | 5 mL 10% (v/v) TritonX-100 |
| 2mM EDTA pH 8.0 | 200 µL 0.5M EDTA pH 8.0 |
| 20mM Tris-HCl, pH 8.0 | 1 mL 1M Tris-HCl, pH 8.0 |
| ddH2O | ddH2O to volume |

Table A6. High salt wash buffer

| **High salt wash buffer (4°C)** | **50 mL High salt wash buffer** |
| --- | --- |
| 500mM NaCl | 5 mL 5M NaCl |
| 0.1% SDS | 0.5 mL 10% (w/v) SDS |
| 1% TritonX-100 | 5 mL 10% (v/v) TritonX-100 |
| 2mM EDTA | 200 µL 0.5M EDTA |
| 20mM Tris-HCl, pH 8.0 | 1 mL 1M Tris-HCl, pH 8.0 |
| ddH2O | ddH2O to volume |

Table A7. Lithium Chloride buffer

| **LiCl buffer (4°C)** | **50 mL LiCl buffer solution** |
| --- | --- |
| 0.25M LiCl | 3.125 mL 4M LiCl |
| 1% (v/v) NP-40 | 2.5 mL 20% (v/v) NP-40 (Tergitol™ 70% in H2O Sigma-Aldrich NP40S-100ML) |
| 1% Sodium Deoxycholate | 0.5 g sodium deoxycholate |
| 1mM EDTA pH 8.0 | 100 µL 0.5M EDTA pH 8.0 |
| 10mM Tris-HCl, pH 8.0 | 0.5 mL 1M Tris-HCl, pH 8.0 |
| ddH2O | ddH2O to volume |

Table A8. TE buffer

| **TE buffer (4°C)** | **50 mL TE buffer solution** |
| --- | --- |
| 10mM Tris-HCl, pH 8.0 | 0.5 mL 1M Tris-HCl, pH 8.0 |
| 1mM EDTA pH 8.0 | 100 µL 0.5M EDTA pH 8.0 |
| ddH2O | ddH2O to volume |

Table A9. SDS elution buffer

| **SDS Elution buffer (RT)** | **For 10 mL** |
| --- | --- |
| 1% SDS | 1 mL 10% (w/v) SDS |
| 0.1M NaHCO_3_ | 84 mg NaHCO_3_ |
| ddH_2_O | Make to volume ddH_2_O |
